# Offline cerebello-cortico-striatal dynamics predict motor strategy exploration and retention in skill learning

**DOI:** 10.64898/2025.12.09.693142

**Authors:** Romain W Sala, Ahsan Ayyaz, Clément Léna, Daniela Popa

**Author notes:** Clément Léna and Daniela Popa jointly directed the work. Corresponding Authors: Daniela POPA, IBENS, 46 rue d’ Ulm, 75005 Paris. Clément LÉNA, IBENS, 46 rue d’ Ulm, 75005 Paris.

## Abstract

Learning a motor skill requires exploring multiple possible solutions to find and retain an optimal strategy^1–3^. The exploration and consolidation of the motor strategy are usually considered to be segregated respectively between task practice and rest periods^4–7^. Here we show that two types of offline reactivations of neuronal representations of locomotor strategies in a brain-wide motor circuit are associated with exploration and retention. During short rests between trials, replays of these brainwide representations predict subsequent shifts in strategies indicating that distributed offline processes participate in strategy exploration. Shifts in strategy are consolidated, and their retention follows population reactivation in cerebellar sleep spindles and stabilization of cerebello-cerebral functional connectivity patterns, consistent with the role of spindles in brain plasticity. Overall, motor strategy optimization and consolidation are supported by two intertwined but distinct types of offline distributed reactivation events involving interregional interactions.

## INTRODUCTION

Motor skill acquisition is a complex process that improves performance through extensive repetition ^1,3^ and error reduction ^8^, ultimately leading to an optimal solution ^2^. How the brain navigates and retains a strategy among multiple possible options is a central question to understand motor learning. While skill learning reduces motor variability, which may arise from both internal and external factors ^9–13^, recent studies suggest that variability itself may aid learning by allowing exploration of new motor solutions ^11,14–16^. This variability is indeed adaptively influenced by past experiences and tasks ^16–18^, facilitating flexible exploration of movement strategies ^6^.

Brain imaging reveals that skill learning is accompanied by reorganizations in motor-related circuits, notably the cerebral cortex, cerebellum and basal ganglia ^19^. These structures exhibit distinct learning capacities ^20^ which are nonetheless interdependent, as shown by cerebellum-dependent skills requiring functional basal ganglia for proper consolidation ^21,22^. Neural reorganization of motor brain circuits can occur during rest between practice sessions, further enhancing motor performance and retention ^23–25^. Indeed, it has long been known that resting periods, even on the scale of tens of seconds or minutes, interspersed between repetitions of the task (“distributed” practice) may be more efficient for task learning compared to uninterrupted repetitions (“massed” practice) ^7^, an effect usually attributed to consolidation ^26,27^. Additionally, sleep may favor consolidation, notably via sleep spindles ^28^, which are brain activity patterns occurring in non-REM sleep and are associated with memory strengthening ^29^.

In this study, we examined how the offline activity of the main brain motor regions relates to the learning of skilled locomotor strategies in an accelerating rotarod task. In this task, mice must stay on a horizontally rotating rod that gradually speeds up; mice are trained in daily sessions of multiple trials interspaced by rest periods lasting a few minutes. The task recruits a motor network involving the motor cortex, basal ganglia, and cerebellum ^30–35^. We found that to stay on the accelerating rod, mice use a locomotor strategy composed of a mixture of gaits, which is adjusted over multiple trials and retained across training days when mice are allowed to pause for several minutes between successive trials but not otherwise. Each gait type corresponds to specific neuronal activities within the motor network. These distributed activities also occur spontaneously during rest between trials and predict intertrial changes in strategy. Learning is associated with a stronger functional connectivity (i.e. statistical interdependence of remote activities^36,37^) within the network observed during the motor execution, and offline during sleep spindles that involve the cerebellum. Task-related reactivations during these spindles appear to support the consolidation of locomotion strategies.

### Changes of motor strategy along learning

Progress in accelerating rotarod performance results in an increase in latency to fall (n=5 mice, Fig. 1ab). However, this classical metric of motor improvement provides little information on how this progress is achieved. Increases in latency to fall have indeed been reported to be accompanied by changes in locomotor strategy along learning ^38^ as well as improved interlimb coordination ^39^.

**Figure 1.**
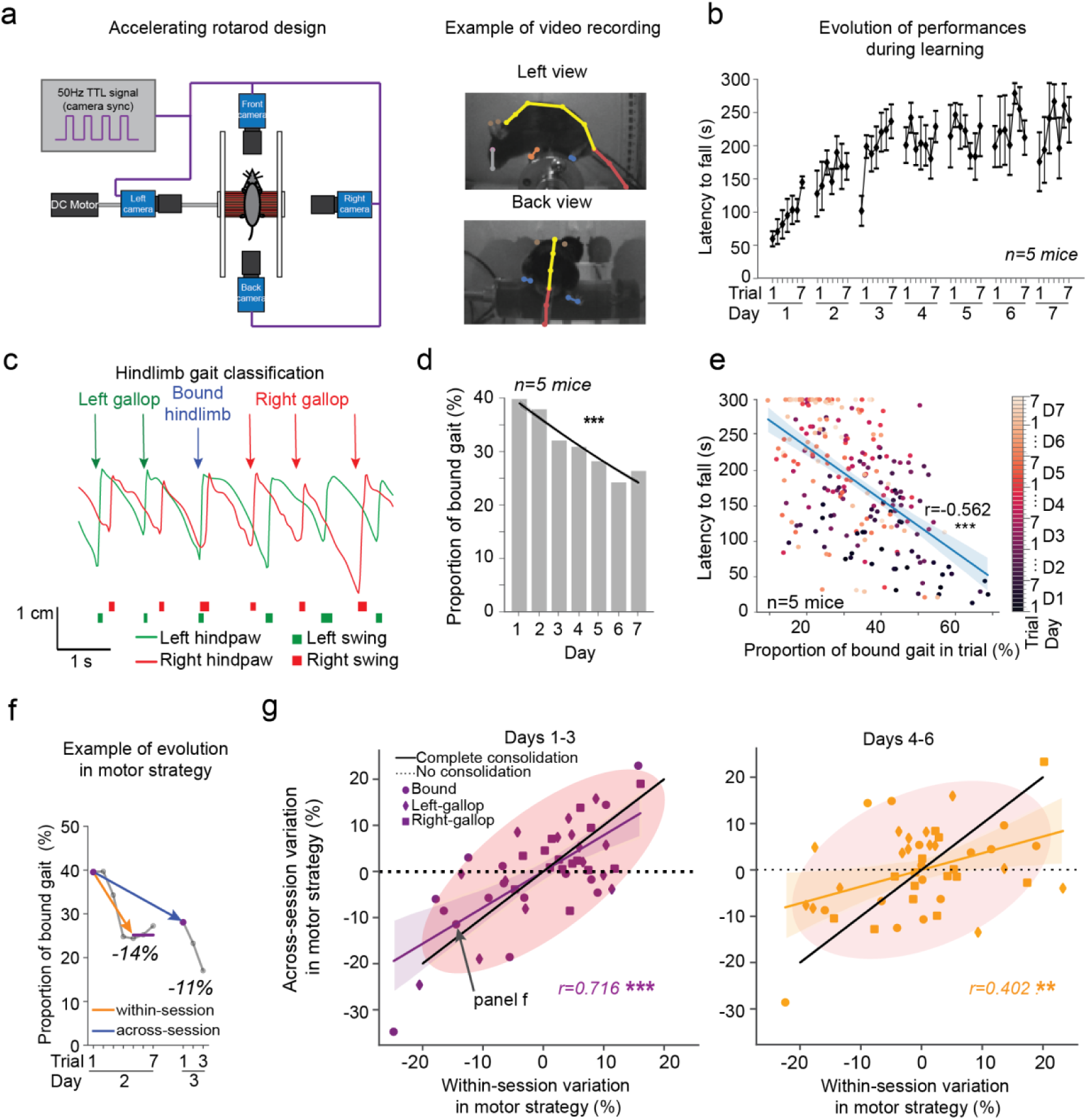
Accelerating rotarod learning is associated to within-session exploration of strategy and across-session retention of changes in motor strategy. **a)** Schematic of the accelerating rotarod setup (left), frame examples with limb and body landmarks from automated tracking; left view and back view cameras (right). **b)** Latency to fall during the accelerating rotarod across trials along days in 5 mice recorded over the 7 days protocol (7 trials/session). **c)** Example traces for the vertical positions of the hind-paws in time, as well as the associated hindlimb gait classification. Colored arrows represent the different gait types. Left and right swings are reported below the traces. **d)** Evolution of the hindlimb bound gait usage along learning days. Logistic regressions are represented. ***p<0.001, Chi-square test on logistic regression. **e)** Scatterplot and associated linear regression showing the latency to fall as a function of the percentage of bound gait usage during trial. The colors of the dots reflect the trial number. ***p<0.001, Pearson’s correlation test. **f)** Example of evolution of bound gait usage showing a within-session decrease between early trials (first trial) and late trials (3 last trials), and an across-session decrease between early trials of consecutive sessions. **g)** Scatterplot comparing within-sessions and across-session variations in motor strategy, for days 1 to 3 (left, across session to days 2 to 4) and days 4 to 6 (right, across session to days 5 to 7), revealing a more efficient consolidation of changes in motor strategy during early learning. Covariance ellipses of 2s.d. and linear regressions are represented. ***p<0.001, Pearson’s correlation test. See supplementary tables for details on samples and statistics.

To document the evolution of motor strategies used throughout rotarod learning, we built a custom rotarod with transparent walls allowing us to track hindlimb movements during the task (Fig. 1a). Examining the kinematics of the hindpaws (Fig. 1c) revealed that mice used a combination of gaits on the rotarod: in many steps, both hindlimbs moved together (overlapping swing phases of both limb), as in the ‘bound gait’ on ground locomotion ^40^. Alternatively, mice used asymmetric hindlimb patterns with a relative phase of hindlimb swing onset close to 90° phase shift (Fig. 1Sup1a, Fig. 1Sup2a). This is reminiscent of the relative hindlimb phase in gallop on ground locomotion ^40^ so we refer to these gait patterns as ‘hindlimb left-gallop’ if the swing of the left hindpaw started just before the swing of the right hindpaw, and ‘hindlimb right-gallop’ otherwise (Fig. 1c).

However, while bound gait and gallop in ground locomotion typically involve uninterrupted sequences of steps at high frequencies (about 10 steps per second) corresponding to locomotion speeds close to 100 cm/s ^40^, steps on the rotarod occurred at much lower frequencies (from 1 to 2 steps per second), consistent with the rotarod moving at lower speeds below the mouse body (from 1.6 cm/s to 7.1 cm/s, Fig. 1c). Thus, although we use the terminology used in ground locomotion to refer to the gaits on rotarod, the accelerating rotarod actually requires the adoption of specific locomotor strategies, which only bear partial resemblance to those found in ground locomotion.

The proportion of bound gait expressed by the mice on the rotarod decreased along learning down to about ~25% (Fig. 1d), and was anti-correlated to the latency to fall (Fig. 1e), suggesting that high proportion of bound gait is not an effective motor strategy on the rotarod. This change is robust, as it is visible at every speed regime, except for the highest rotarod speed (>40rpm) (Fig. 1Sup1a), at which the proportion of bound gait is already low from the first learning sessions.

Strikingly, a gradual change in gait preference takes place across the multiple trials in learning sessions. We then examined if these within-session variations in gait preference were maintained overnight into the next session. Indeed, the variations of gaits usage in trials between the beginning and the end of one session (‘within-session’, Fig. 1f) were strongly correlated to the variations between the beginning of one session and the beginning of the next session (‘across-session’, Fig. 1f) for the first three learning days as evidenced by the correlation between within- and across-sessions variations; it became weaker on later days (Fig. 1g). Although the retention across-session of changes in motor strategy was overall more prominent during earlier days compared to later days (Fig. 1g), we did not observe a monotonous decrease in consolidation along the 7 days of training (Fig. 1Sup1c).

We then sought to verify that the adjustment of motor strategy, as defined as gaits usage, was indeed the dominant learnt feature visible during accelerating rotarod learning, rather than a secondary consequence of improved inter-limb coordination or limb kinematics. Thus, we examined the evolution of these parameters for each type of gait. Analysis of fore-hind limb coordination revealed that the forelimbs were overall synchronous with their homolateral hindlimb (Fig. 1Sup2a). While each gait exhibited characteristic features of inter-limb coordination (Fig. 1Sup2ab), we did not observe major within-gait changes in coordination along learning neither for the relative phases nor the variability of the relative phases (resp. circular mean and circular standard deviation, Fig. 1Sup2b). Similarly, the fore-and hind-limb kinematics were stable along learning sessions (Fig. 1Sup2c). Overall, these results indicate that although the motor strategy used during the task evolves consistently along learning, the underlying features of each type of gait —interlimb coordination and limb kinematics— remained comparatively invariant.

We then examined to which extent the within-sessions adjustments of motor strategy, and their maintenance across sessions, involve offline processes occurring outside the task, particularly during the resting periods in-between the trials. To this end, we subjected two groups of mice to the accelerating rotarod learning, one control group with 5-min intertrial resting periods (n=5 mice), and one with short resting periods (n=5 mice, intertrials’ median duration of 55s and median absolute deviation of 9s), thus limiting the amount of offline neural processes potentially occurring in-between trials (Fig. 1Sup3a). Mice exposed only to short resting periods displayed lower levels of learning compared to control animals (Fig. 1Sup3a), and a marked reduction in session-to-session retention of performances (Fig. 1Sup3b), indicating an impaired consolidation of shifts in motor strategies across sessions. Whereas control mice exhibited a robust within-session reduction in bound gait which was maintained across-session (Fig. 1Sup3cde), mice subjected to short rest showed only shallow and inconsistent within-session changes in motor strategy (Fig. 1Sup3de), and within-session changes were less reliably maintained into the next session (Fig. 1Sup3fg). These results collectively indicate that truncating intertrial rest periods disrupts the offline refinement and consolidation of motor strategy that normally support rotarod learning.

Overall, these observations indicates that the accelerating rotarod learning is associated with a progressive refinement of the motor strategy used by the mice across trials, which is efficiently carried to the next session (particularly during the first days of learning). In addition, these processes seem to require resting periods between trials, which is consistent with the existence of an offline consolidation process in accelerating rotarod learning ^35^.

We then used this paradigm to investigate the brain neurophysiological activities associated with the refinement of motor strategy and to its consolidation.

### Recording cerebello-cerebral pathways

We simultaneously recorded the neuronal activitiy of single units in the cerebellum, basal ganglia and motor cortex, to study their coordination during the learning and consolidation of changes in the motor strategy in the accelerating rotarod. We therefore implanted bundles of electrodes in the left cerebellar nuclei (CN, including both Dentate and Interposed nuclei), bilaterally in the hindlimb primary motor cortex M1 and dorsolateral striatum DLS of mice (n=5, Fig. 2a). We also recorded from the right motor thalamus (ventral-anterior lateral thalamus, VAL) and centro-lateral nucleus of the thalamus (CL) which relay the left CN inputs to M1 and DLS ^35^. Overall, we recorded 1076 units over the 7 days of rotarod training of 5 mice (see Supplementary table 1 and Methods section of details).

**Figure 2.**
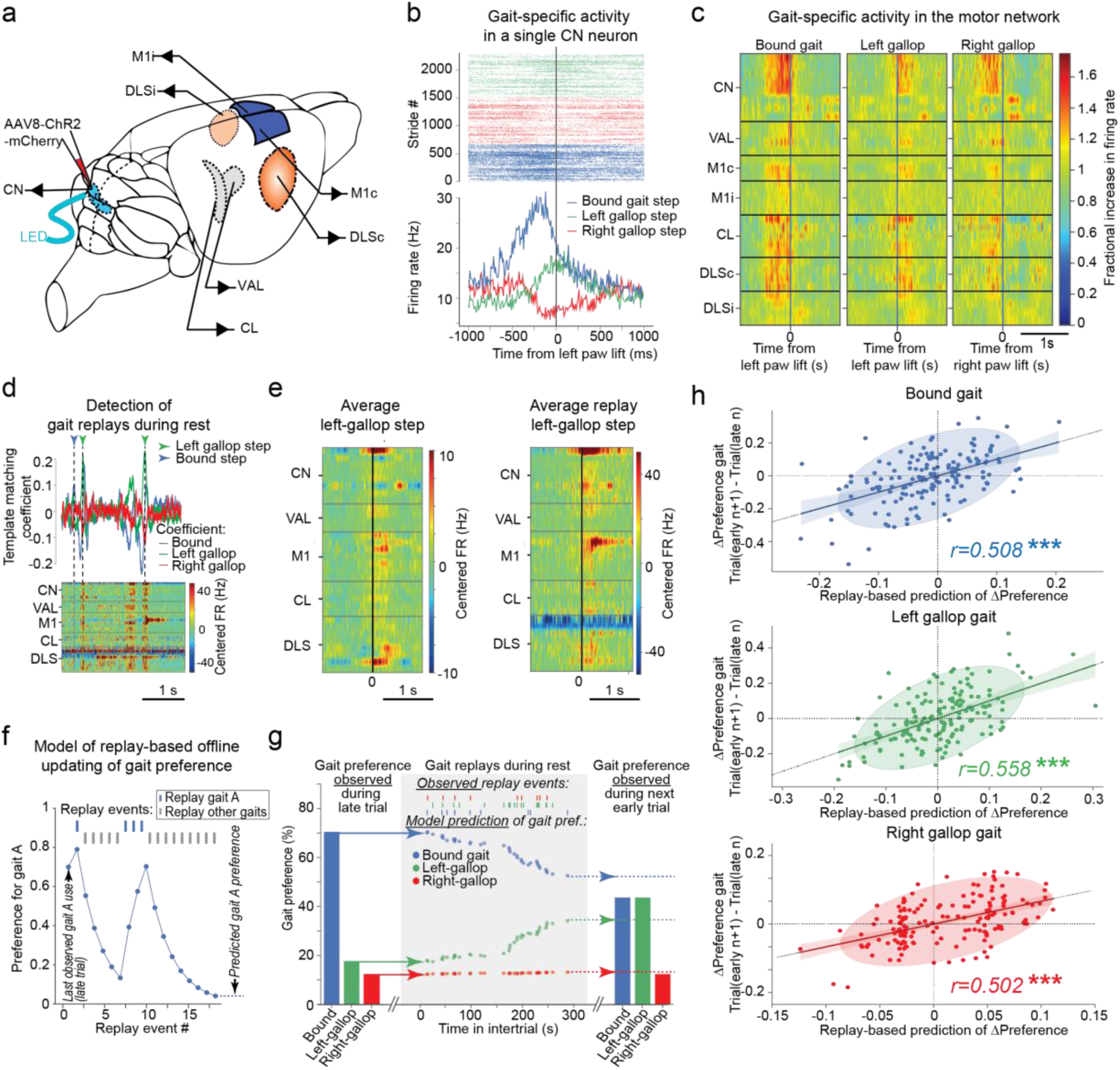
Strategy-specific replays of neuronal activity in the motor network during resting are predictive of inter-trial changes in motor strategy. **a)** Experimental design for the extracellular recording with bundles of electrodes in the cerebellar nuclei, thalamus, motor cortex and striatum, and viral strategy for the optogenetic activation of the cerebellar nuclei. **b)** Example of raster plot and peri-stimulus time histogram (PSTH, 10ms bins) centered on the lift of the left paw for a CN neuron during the different types of gaits. **c)** Example of heatmap representation of PSTHs, centered on paw-lift for neurons recorded simultaneously in a same mouse in the CN, VAL, contra- and ipsilateral M1 (M1c, M1i), CL, contra- and ipsilateral DLS (DLSc, DLSi) during the different types of gaits. **d)** Example of template matching analysis (top) detecting the replays of different types of gait (vertical lines) from the neuronal activity of a mouse during resting period (bottom). Replays are considered as significant when the correlation with the template exceeds 5 s.d. above the correlation with shuffled traces **e)** Example of strategy-specific patterns of activity replayed during resting periods and observed during task (M1 and DLS are each pooled for ipsi- and contralateral electrodes). **f)** Schematic learning model used to predict trial-to-trial change in gait proportion based on the temporal structure of the replay during a resting period. **g)** Example of gait proportion in two subsequent trials (late trial on the left and next early trial on the right) and learning model predictions for the trial-to-trial change in gait proportion based on the temporal structure of the replay during a resting period (center). **h)** Scatterplots and associated linear regressions showing the actual variation in gait proportion between trial-to-trial as a function of the predicted variation for different for different gait types. Covariance ellipses of 2s.d. are represented. ***p<0.001, Pearson’s correlation test. 1076 cells from 5 mice, 7 days. See supplementary tables for details on samples and statistics.

In these mice, CN neurons were infected with an AAV8-hSyn-hChR2(H134R)-mCherry, and an optical fiber was positioned above the CN. 100ms optogenetic stimulations of the CN during open-field sessions triggered short latency responses of neurons in VAL and M1, with significantly shorter latencies in VAL (Fig. 2Sup1), consistent with the positioning of the electrodes along a cerebello-thalamo-cortical pathway. Similarly, short latency responses were observed in CL and DLS, with shorter latencies in CL (Fig. 2Sup1) consistent with the positioning of these electrodes along a cerebello-thalamo-striatal pathway.

### Replays of gait-specific activities

We then examined the neuronal modulation across brain regions during accelerating rotarod learning. Gait-related modulations were observed throughout the recorded regions: “bound gait”, “left-gallop” and “right-gallop” types of hindlimb gaits were associated with distinct modulations in terms of magnitude, polarity and temporal sequence across the network (Fig. 2bc). In each structure, the neurons generally displayed low correlations between their patterns of modulation during the different gait types (absolute median below 0.2, Fig. 2Sup3a), thus indicating gait-specificity was present throughout the network.

These activity patterns could reflect incidental and independent recruitment of neurons in each structure by sensorimotor processes, or alternatively arise because these regions belong to a strongly interconnected network, in which case coordinated activity could also emerge outside the actual execution of the task. Thus, we examined the occurrence of the coordinated “offline” reactivations of these neuronal assemblies across brain structures during the rest periods between the trials in the absence of motor activity. We used a template matching approach ^41–43^ validated by activity shuffling to detect the presence of gait-like replay events displaying similarity to the sequence of activity induced across brain regions by each type of gait (Fig. 2de). Strikingly, this approach revealed intermittent (~0.1-0.2 Hz, Fig 2Sup4c) “replay” events, involving a coordinated activation of neurons in the cerebellum, thalamus, motor cortex and striatum corresponding to different types of gaits (Fig. 2Sup4bc). These events revealed to be specific to the type of gait that they were attributed to, as evidenced by the low values of disambiguation coefficients (Fig. 2Sup4d), suggesting independent reactivations corresponding to the different strategies used during the accelerating rotarod.

We then evaluated the relative preservation of sequential neuronal firing within these replay events, and compared it to the consistency observed between individual gaits. For this, we implemented the matching index metric (Fig. 2Sup4e, see methods), which revealed that the detected replay events displayed slightly higher levels of consistency with gaits or within themselves (Fig. 2Sup4ef) than the levels of consistency observed within gaits (Fig. 2Sup4g). This correspondence between offline and online activity across multiple motor regions suggests a tight functional interdependence of neuronal firing between these structures beyond motor actions.

### Replays predict motor strategy changes

Gait replay events during immobility episodes were not specific to the rotarod since they could be observed in the open-field preceding the rotarod sessions (Fig. 2Sup5a), indicating that our recordings could not distinguish these events from pre-existing gait patterns likely originating from ground locomotion. However, we observed that the increases or decreases in proportions of gait replay events during intertrial rest compared to the open-field session before the rotarod training were preserved in the open-field session following it (Fig. 2Sup5bc), suggesting a link between these proportions and task learning.

We then investigated whether the proportion of replays during resting periods could be linked to trial-to-trial changes in motor strategy. To account for adjustments of strategy, which occur during trials, we only considered the late part of the trial preceding the resting period (second half of the trial) to be the reflection of the strategy used at the end of the trial, before the resting period. And conversely, we only considered the early part of the subsequent trial (first half of the trial) to reflect the initial strategy used after the resting period.

Since offline activity has also been linked to consolidation of learning in other systems (e.g., ^22,44–47^), we first examined if the proportions of the different gait-like offline activity recapitulate the proportion of gaits used before the rest period. We only observed a weak anticorrelation in the case of the right gallop, but neither for bound nor for left gallop (Fig. 2Sup6a). Alternatively, gait-like offline activity during rest between trials could directly predict the initial combination of gaits used after the resting period. However, we only observed a weak anticorrelation in the case of the right gallop, but neither for bound nor for left gallop (Fig. 2Sup6b). This indicates that the proportion of the different offline gait-like activities during the intertrial epochs did neither recapitulate nor predict the proportion of types of gaits used in the previous and next trials.

We then hypothesized that the intertrial activity was predicting the shift in locomotor strategy between successive trials rather than the strategy itself. Indeed, a higher (or lower) proportion of intertrial replays of a given gait relative to the use of this gait in the previous trial predicted a corresponding increase (or decrease) in the use of that gait in the next trial relative to the previous trial, as indicated by the positive correlations between these relative proportions (Fig. 2Sup6c). To more thoroughly test this hypothesis, we defined a simple learning model of “gait preference”, a parameter determining the probability of using a given gait during the behavior. According to our model, the update of this preference during intertrial resting periods is governed by inter-trial gait-like offline events (Fig. 2fg). In the model, each offline occurrence of a given gait-like activity (e.g., bound-gait like activity) increases the preference for this type of gait, while the offline occurrence of another gait-like activity will decrease it. This simple model only requires determining a ‘learning rate’ parameter which defines the fraction of gait-preference updated at each replay event for all sessions of a mouse, a “gain” parameter (scaling the offline update into an expressed shift in preference between successive trials), and an “offset” parameter (which defines the relative rate of replay yielding a constant gait preference). Strikingly, we found strong significant correlations between the change in gait preferences predicted by this simple model of intertrial updating of gait preference and the actual change in gait preference observed between successive trials (Fig. 2h).

We also verified that training and evaluating these models on randomized intertrial variations of motor strategy (Fig. 2Sup7a), or on precedent and subsequent intertrial variations of motor strategy (Fig. 2Sup7b), generally yielded models with lower performances. This supports the correspondence between “replays” in a given intertrial epoch and the corresponding changes in the motor strategy observed in the next trial.

Altogether, these results suggest that offline gait-like events may influence the refinement of the motor strategy in a trial-to-trial basis by adjusting the preference for each type of gait.

### Increased cerebello-cerebral coupling

The observed coordination of firing across neurons of the motor circuits could be refined as locomotor strategies improve ^38^. Since these brain regions are connected by oligosynaptic pathways, we first examined the presence of sequential activation of neurons located in different structures. We therefore computed the cross-correlograms of spike trains from neurons in the CN, M1 and DLS during each rotarod trial. While these cross-correlograms generally exhibited a broad central peak consistent with a covariance of firing rate of the pairs of neurons, they also revealed, in some pairs, an enrichment of spiking of one neuron shortly after the firing of the other, as revealed by asymmetric correlograms (Fig3a). Subtracting the left-side of the cross-correlogram from its right side revealed the temporal structure of this asymmetry, which in average peaked below 25 ms (Fig. 3Sup1ab). Such patterns of sequential activation are hereafter referred to as directional functional coupling between CN, M1 and DLS.

**Figure 3.**
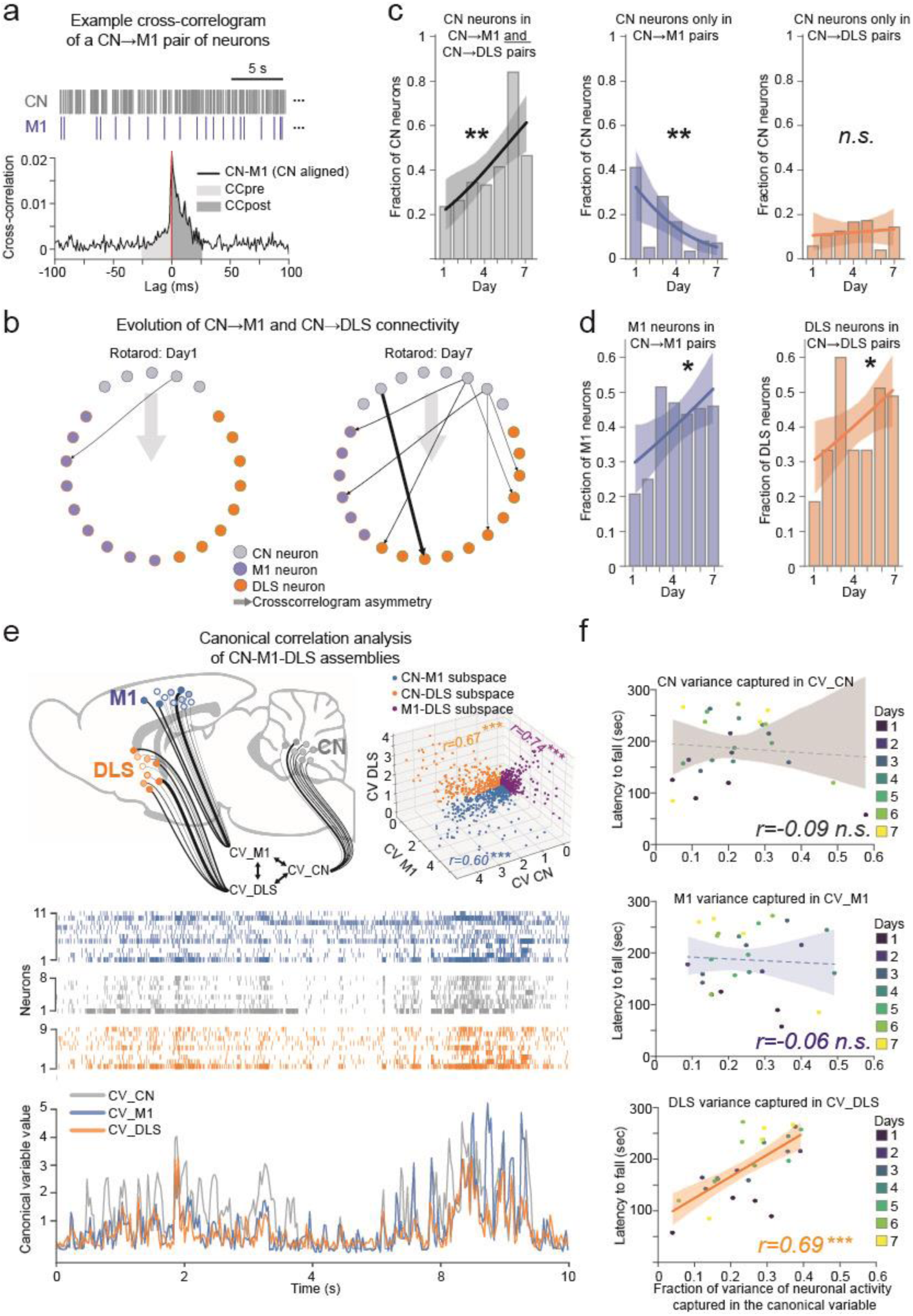
Accelerating rotarod learning is associated to an enhanced coordination of the cerebellum, the motor cortex and the striatum during the task. **a**) Example cross-correlogram of a CN neuron and a M1 neuron and associated spiketrains (top). **b)** Summary of pairs with asymmetric correlations observed from single sessions of a single mouse represented as a connectivity map of CN, M1 and DLS neurons displaying significant CN→M1 and CN→DLS connectivity for a mouse during the first day (left) and during the last day of rotarod training (right). Significant connectivity during trials is represented by arrows with a thickness proportional to the directional index. **c)** Evolution of the fraction of CN neurons exhibiting significant outbound interaction with neurons in both M1 and DLS (left), only in M1 (middle), only in DLS (right), and along learning days. Logistic regressions are represented. **p<0.01, Chi-square test on logistic regression. **d)** Evolution of the fraction of M1 neurons (left) and DLS neurons (right) exhibiting significant inbound interaction from CN neurons along learning days. Logistic regressions are represented. *p<0.05, Chi-square test on logistic regression. **e)** Example of multiview canonical correlation analysis (MCCA) revealing cerebello-cortico-striatal assemblies during accelerating rotarod (top-left). The thickness of the arrows is proportional to the weight of neurons. Example rasterplot (center) and associated canonical variables (bottom). 2-dimensional projections of the canonical variables in CN-M1, CN-DLS and M1-DLS subspaces (top-right). **f)** Scatterplot comparing the fraction of CN (top), M1 (center) and DLS (bottom) activity variance captured by MCCA to the latency to fall, revealing a correlation between the fraction of striatal activity captured by the MCCA and the average performance of mice. ***p<0.001, Pearson’s correlation test. See supplementary tables for details on samples and statistics.

We then examined whether the directional functional coupling between regions evolved along learning. We therefore identified, for each day of training, pairs of neurons that showed significantly asymmetric cross-correlograms. In such pairs, the spiking of single CN neurons could be followed either by increased firing probability (within 25 ms) of M1 neurons only, DLS neurons only, or both (Fig. 3b). Strikingly, we found that over the 7 days of training, the fraction of our sample of CN neurons functionally coupled with both M1 and DLS neurons increased (Fig. 3bc), while the fraction of CN neurons exclusively engaged with M1 decreased (Fig. 3bc), suggesting that the CN initially engages preferentially with M1, but that the accelerating rotarod learning promotes coordinated interactions of CN neurons with DLS and M1 neurons. In addition, the fraction of M1 and DLS neurons firing after CN neurons increased over the 7 learning sessions (Fig. 3bd), signaling that accelerating rotarod learning induces an increase in directional functional coupling of the cerebellum with both M1 and DLS.

We also assessed the intensity of directional functional coupling between neurons by building a “directional coupling index” quantifying the asymmetry of cross-correlogram. Interestingly, this index did not reveal substantial changes in the intensity of coupling in pairs of cells from each pair of structures during rotarod execution (Fig. 4acd), signaling that the increase in functional interactions between structures is primarily associated with an increase in the number of neurons in the target regions entrained by the cerebellum.

### Task-related population activity

The increased functional coupling of the cerebellum with the motor cortex and the striatum may result in the emergence of coordinated cerebello-cortico-striatal assemblies of neurons. To identify such assemblies of co-varying neurons in the network, we used a Multiview Canonical Correlation Analysis (MCCA). This analysis identifies assemblies of neurons in each region which display maximally correlated activities (Fig. 3e) and thus provides an integrated measure of the coordination of neuronal activity across these regions during the task. The activity of these assemblies is composed of a weighted sum of neuronal firing rates in each region (called “canonical variable”, or CV), where the weight of individual neurons in this linear combination reflects the contribution of the individual neuron to the assemblies’ activity.

We thus performed MCCA on trials using neurons recorded in CN, M1 and DLS (Fig. 3e). This revealed the existence of a first MCCA component, typically explaining ~25% of the variance in each region (Fig. 3Sup2a) and exhibiting high correlations (>0.6; Fig. 3Sup2bc) between the canonical variables of the three structures, consistent with the recruitment of cerebello-cortico-striatal assemblies displaying a coordinated activity during the task. Strikingly, the fraction of the activity of the DLS captured by MCCA was directly correlated with the performance of mice during each session (Fig. 3f), suggesting that an efficient learning is associated with an increased participation of the DLS to cerebello-cortico-striatal assemblies. Interestingly, the fraction of the activity of the CN and M1 captured by the MCCA wasn’t correlated to the performances (Fig. 3f), supporting the idea that motor learning involves the coordination of the DLS with the cerebello-cortical network.

### Reactivation of task-related assemblies

To assess the extent to which the inter-regional directional functional coupling observed during the task is re-expressed in offline spontaneous activity ^48^, we first examined the directional coupling of pairs of cells during resting periods along learning. Strikingly, we found that pairs of CN->and M1 or CN->DLS neurons with positive directional coupling index (i.e., M1 or DLS neurons exhibiting increased firing probability within 25 ms after CN firing) during the task also showed positive coupling during resting periods, and the proportion of pairs maintaining significant sequential activation during rest progressively increased along the task (Fig. 4a-d). Indeed, for pairs of neurons showing directional coupling during the task, the directional coupling index during rest increased along learning days, reaching values similar to those observed during the task in the later days of learning (Fig. 4cd). These observations support the idea that task learning is associated with a long-term reinforcement of the directional functional connectivity of networks engaged during the task.

**Figure 4.**
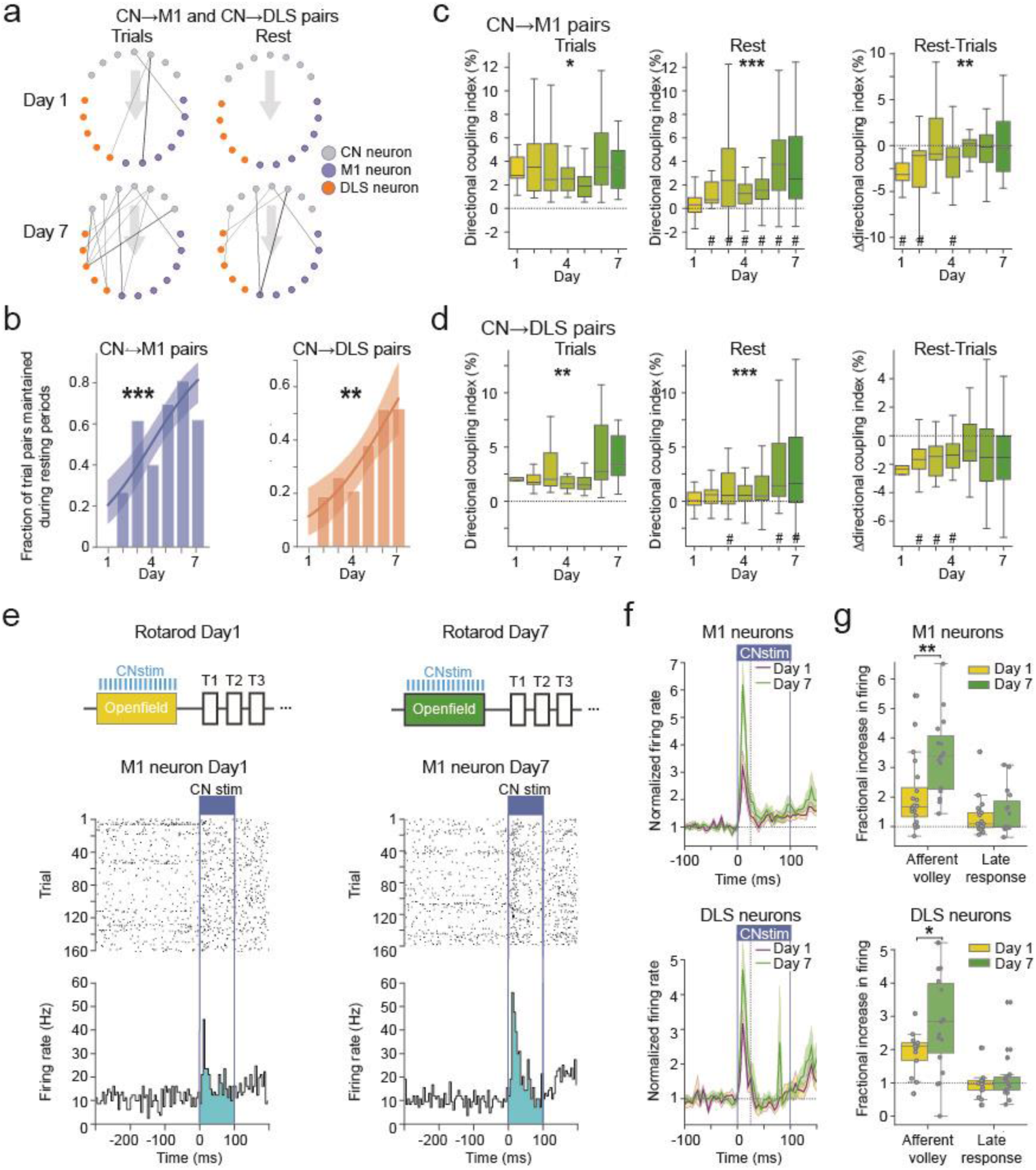
Accelerating rotarod learning induces a long-lasting increase in functional connectivity of the cerebellum with the motor cortex and the striatum. **a**) Summary of pairs with asymmetric correlations observed from single sessions of the same mouse represented as connectivity maps during trials (left) and rest (right) for CN->M1 and CN->DLS pairs, in a mouse during the first (top) and last day of rotarod training (bottom). Significant connectivity is represented by arrows and line width is proportional to the directional index. **b)** Fraction of CN->M1 (left) and CN->DLS pairs significant during trials and maintained during rest along learning days. Logistic regressions are represented. **p<0.01, ***p<0.001, Chi-square test on logistic regression. **c)** Directional indices of CN->M1 pairs significant during trials (left), the same pairs during rest (center) and the difference between trial and rest (right). Outliers are not shown. *p<0.05 **p<0.01, ***p<0.001, Mixed-effect linear model ANOVA. #p<0.05, Wilcoxon test for the difference to 0, Holm-Sidak corrected. **d)** Same as panel b for CN->DLS pairs. **e)** Example of PSTH (5ms bins) and raster plots, centered on CN stimulations onset from M1 neurons of the same mouse during the first day (left) and last day (right) of training. **f)** PSTH (5ms bins) displaying the change in normalized firing rate (normalized to the 300ms before stimulation onset, average ± SEM) of responsive neurons in M1 (top) and DLS (bottom) before the first day and before the last day (vertical dashed lines represent the limit of the afferent volley, 25ms). **g)** Average fractional increase in firing of responsive neurons in M1 (top) and DLS (bottom), during the afferent volley (0 to 25ms) and the late response (25 to 100ms). *p<0.05, **p<0.01, Mann-Whitney U test. See supplementary tables for details on samples and statistics.

Since replays of gait-related activity occur during intertrial rest (Fig. 2), we examined if the sequential activations of neurons took place during these gait-related events. We therefore computed the directional coupling index using neuronal activity only either during episodes of gait-like replays or in time intervals without replays. Reducing the number of spikes used in the analysis decreased the signal-to-noise ratio, but by pooling across all days, we observed positive directional coupling index for CN->M1 pairs both in and out of gait-like replay events and no significant difference in directional coupling index between periods of replays and replay-less rest (Fig. 4Sup1ab). No significant effect was found for CN->DLS pairs, but this may result from the weaker initial effect before the partition in time segments with and without gait-like replays (Fig. 4d). Thus, sequential activation of pairs of neurons observed during the task is increasingly replayed offline along learning and takes place both during and outside of gait-replay events. Therefore, the increase of functional directional connectivity observed during the task does not result directly from the motor activity but instead seems to reflect changes in the organization of the motor circuit, and can therefore be expressed offline between neurons in distant brain structures coupled by oligo-synaptic pathways.

To investigate whether the increases in sequential activations across structures observed along learning during the task and during rest could result from increases in cerebello-cortical and cerebello-striatal oligosynaptic coupling, we examined the impact in M1 and DLS of optogenetic stimulations of the CN neurons. When comparing the optogenetically-evoked responses obtained during openfield sessions of the first (Day 1) and last (Day 7) rotarod training (Fig. 4e), we observed increased responses during a short time window following the stimulation onset (“afferent volley”, first 25ms of stimulation) (Fig. 4fg). In contrast, a similar experiment in a group of mice that was not subjected to accelerating rotarod training but only to daily open field sessions, did not reveal significant increases in response in M1 or DLS1 between Day 1 and Day 7 (Fig. 4Sup2). This is consistent with a potentiation of oligosynaptic pathways linking the cerebellum to the motor cortex and to the striatum.

Altogether, these results confirm that both cerebello-cortical and cerebello-striatal couplings are increased as a consequence of accelerating rotarod learning.

### Cerebellar spindles and connectivity

We then investigated if the presence of task-related reactivations during rest may contribute to the consolidation of learning. For this purpose, we studied sleep spindles, brief oscillatory events generated during slow-wave-sleep in the thalamo-cortical circuit^49^, which are strongly associated with memory consolidation of many forms of learning ^28,50^. Motor learning has indeed been extensively linked to sleep spindles ^51–56^. Moreover, their role may not be limited to the thalamo-cortical system, since they have been also associated with changes in cortico-striatal functional connectivity ^53^. The recent discovery of sleep spindles in the cerebellar nuclei ^57–59^ raise the possibility that these processes participate in changes in connectivity observed in the cerebello-forebrain circuits reported here.

To isolate spindles, we first identified vigilance states of mice using contralateral M1 Local Field Potential (LFP) and observed episodes of sleep^60^, then detected cerebellar spindle-like activity in CN LFP during slow-wave cortical state (Fig. 5a, Fig 2Sup2). These spindle-like events detected in the cerebellum (“CN spindles”) not only entrained neuronal activity in the CN (Fig. 5ab), but M1 and DLS activity was also bilaterally phase-locked to these events (Fig. 5ab). CN spindles were in great majority distinct from spindle events detected in M1c LFP, as only 4.5% of CN spindles were coincidentally associated with the detection of an event in M1c LFP (Fig. 5Sup1abc). In addition, we observed that neurons in the motor network were significantly entrained by either CN or M1 spindle-like events, with only a small fraction of them being locked to both types of events simultaneously (Fig. 5Sup1d). The observed number of neurons locked to both CN and M1 spindle-like events was not significantly lower or higher than expected if being locked to CN and M1 spindles were independent events (binomial tests, see Supplementary Table 42), indicating that neurons in the motor networks are independently recruited by CN or M1 spindles. Overall, these results suggest that most spindles which are detected in the CN and entrain neuronal activity in motor circuits are unlikely to be generated in M1c, but in another upstream thalamocortical-network^61^.

**Figure 5.**
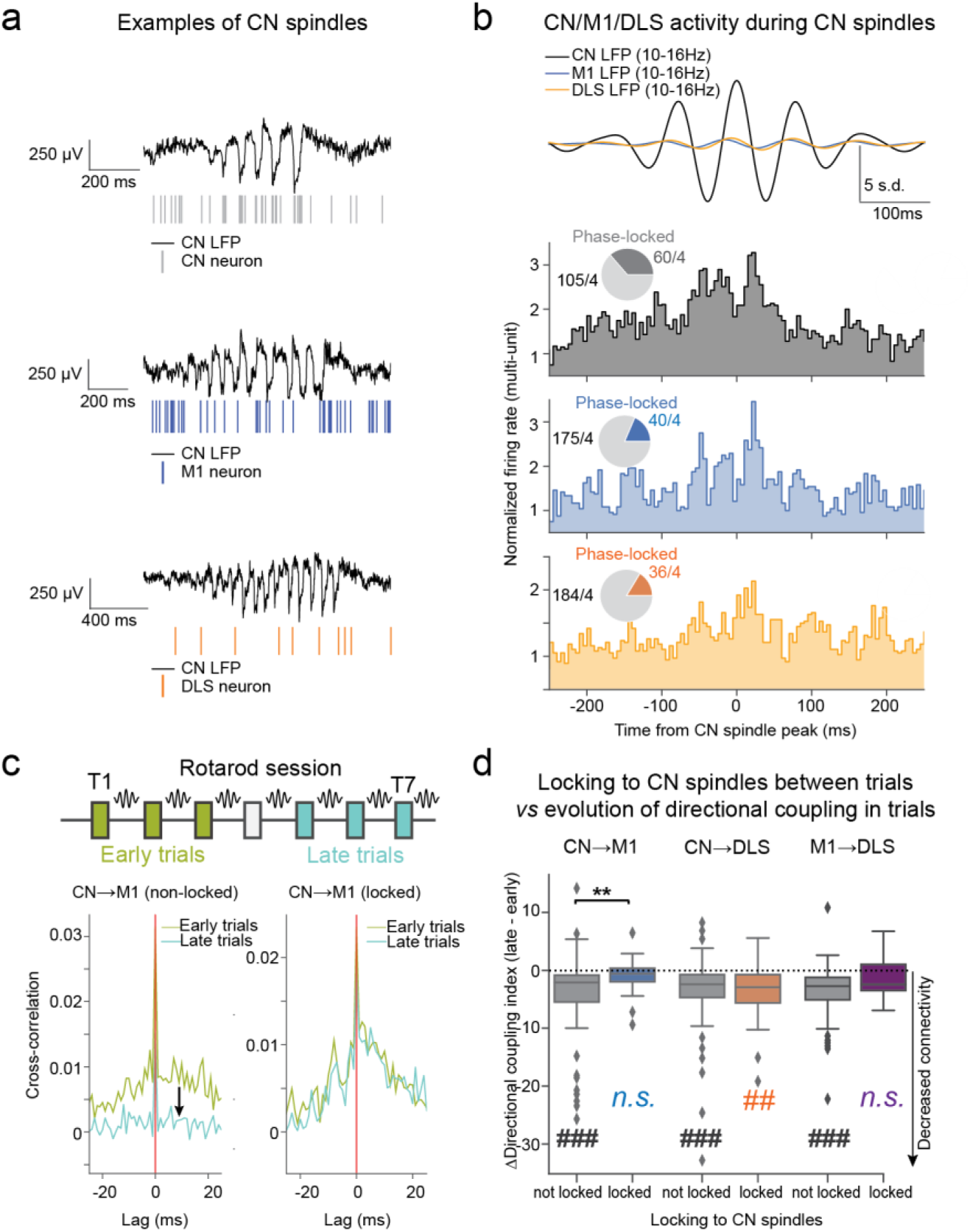
Cerebellar spindles activity during NREM sleep entrains cortical and striatal activity, and is predictive of a maintained functional connectivity in the cerebello-cortico-striatal network. **a)** Examples of spindles detected in the LFP of the CN and simultaneous spiketrains of a CN neuron (top), M1 neuron (center) and DLS neuron (bottom). **b)** Average z-scored filtered LFP (10-16Hz) in the CN, M1 and DLS centered on CN spindles peak (top, a small offset was added to DLS trace for visualization), corresponding normalized multi-unit activity of neurons locked to CN spindles in the CN, M1 and DLS (bottom). Insets correspond to the proportion of significantly locked neurons. **c)** Example cross-correlograms of a CN->M1 pair with the target neuron non-locked to cerebellar spindles (left) and a locked CN->M1 pair (right), respectively displaying a decreased and a maintained functional connectivity during the session **d)** Boxplots showing the evolution of the directional coupling index of interacting pairs during a training session for CN->M1, CN->DLS and M1->DLS pairs depending on their locking of the target neuron to cerebellar spindles. **p<0.01, Mann-Whitney U test. ##p<0.01, ###p<0.001, Wilcoxon test for the difference to 0.

We then examined whether the participation of M1 or DLS neurons to cerebellar spindles during rest between the trials was predictive of changes in their directional functional connectivity with the cerebellum during trials. Strikingly, CN->M1 and M1->DLS pairs with significant directional connectivity during the early trials of the session (i.e., with significant directional coupling index in first three trials) but in which the target M1 or DLS neuron was not entrained by CN spindles displayed a decrease in directional functional connectivity during the late trials of the session (last three) (Fig. 5cd). In contrast, CN->M1 and M1->DLS pairs showing directional functional connectivity in early trials and in which the target neuron was entrained by CN spindles displayed no significant change in directional coupling index (Fig. 5cd). This suggests a role of CN spindles in maintaining the functional connectivity in cerebello-cortico-striatal network. Interestingly, this relationship between locking to CN spindles and persistence in directed connectivity was not visible in CN->DLS pairs, as pairs with DLS neurons either non-locked or locked to CN spindles displayed a decrease in directional coupling index during the session. This indicates that offline expression of task-related cerebello-striatal functional connectivity requires longer timescales to be reinforced and stabilized, in contrast to cerebello-cortical or cortico-striatal connectivity.

Overall, these results suggest that cerebellar spindles are global events entraining cortical and striatal activity bilaterally, which contribute to the stabilization and reinforcement of the functional connectivity in a network linking the cerebellum with the motor cortex and striatum.

### Cerebellar spindles and consolidation

We then examined whether cerebellar spindles could be directly related to the consolidation of accelerating rotarod learning. As described above, task execution is associated with the recruitment of coordinated neuronal assemblies across brain regions captured by MCCA (Fig. 3). We therefore examined the extent to which such brain-wide assemblies were recruited during resting periods. We found that during CN sleep spindles, the covariance of CN, M1 and DLS assemblies was increased compared to eventless resting periods (Fig. 6ab), indicating a reactivation of task-related neuronal populations, which was more likely to occur during CN spindles than outside of sleep spindles or replays. Strikingly, the value of covariance during CN spindles was positively correlated to the retention of performance during the next session (Fig. 6c), suggesting a contribution of CN spindles to the offline consolidation of accelerating rotarod learning.

**Figure 6.**
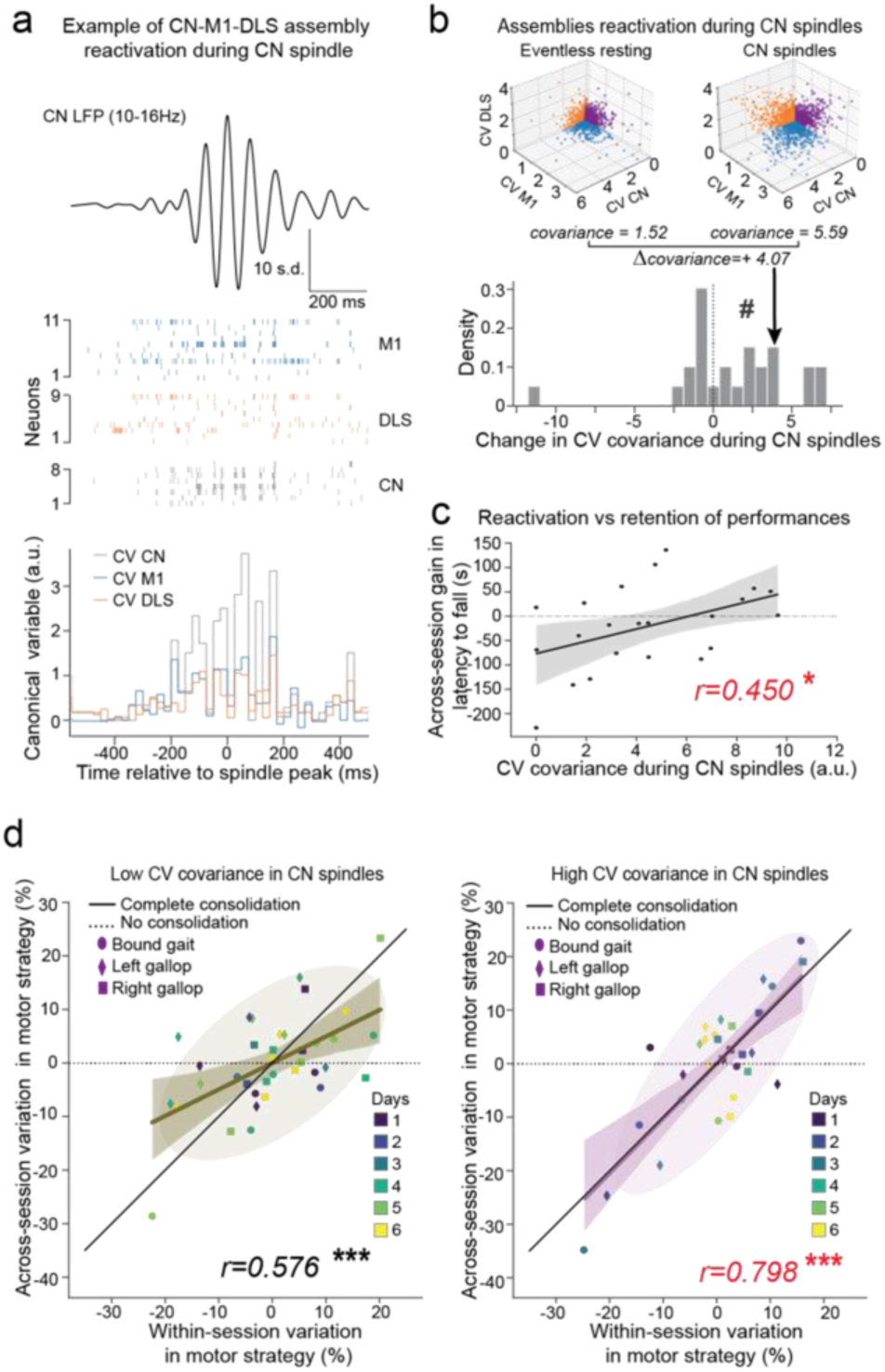
A strong entrainment of cerebello-cortico-striatal assemblies by cerebellar spindles predicts an efficient consolidation of motor strategy and a higher retention of performances. **a)** Example of cerebellar spindle (top), rasterplot showing concomitant neuronal activity in CN, M1 and DLS (center) and associated MCCA canonical variables (bottom). **b)** 2-dimensional projections of the canonical variables in CN-M1, CN-DLS and M1-DLS subspaces of the same mouse and session showed in g, during eventless resting (top-left) and during CN spindles (top-right), displaying an increase in covariance during CN spindles. Histogram showing the increase in covariance during CN spindles (bottom). ###p<0.001, Wilcoxon test. **c)** Scatterplot comparing the gain of latency to fall across-session to the value of covariance during CN spindles. Linear regression is represented. *p<0.05, Pearson’s correlation test. **d)** Scatterplot comparing within-sessions and across-session variations in motor strategy, during sessions with low covariance (left) and high covariance (right) during CN spindles, revealing that a high covariance during CN spindles is associated to an efficient consolidation of changes in motor strategy. Covariance ellipses of 2 s.d. and linear regressions are represented. ***p<0.001, Pearson’s correlation test. See supplementary tables for details on samples and statistics.

Moreover, we also found that sessions that exhibited strong task-related reactivation in spindles showed a clear consolidation across sessions of the changes of motor strategies which took place during the session, while sessions with weaker reactivations in spindles showed less consistent consolidation across sessions. This was evidenced by an anti-correlation between covariance and consolidation error (Fig6Sup1b), as well as an incomplete consolidation (slope <1) during sessions with weaker reactivations (Fig. 6d, 6Sup1c). Although sessions with high covariance were observed preferentially during early phases of learning, we did not observe an overall significantly decrease across sessions (Fig. 6Sup1a). Neither the retention of performances nor the consolidation of motor strategy could be predicted from the session day (Fig. 6Sup1de) indicating that the simple number of repetitions of the task is poor predictor of changes in consolidation, contrarily to the intensity of spindle reactivations. Finally, we observed that, contrarily to reactivations (covariance) during CN spindles, there was no significant correlation of the reactivation during gait replays with the consolidation of performance or with the consolidation of strategies (Fig. 6Sup1fg), which further supports distinct roles of gait replays and spindles in the offline consolidation of accelerating rotarod learning.

Taken altogether, our results suggest that cerebellar sleep spindles, in addition to shaping functional connectivity in the motor network, also contribute to the consolidation of motor learning by allowing the entrainment of brain-wide assemblies of neurons involved in the task.

## DISCUSSION

Our results reveal that motor skill learning is accompanied by increased interactions between the cerebellum, motor cortex and striatum not only during the execution of the learned task but also offline during intertrial periods. We found two types of distinct but intermingled offline interactions, associated with different aspects of motor skill learning. First, the trial-by-trial exploration of locomotor strategies is supported by a differential expression of strategy-specific replays of neuronal activity in the motor network. Second, the long-term retention of the choice of strategy is reflected in brain-scale reactivations during cerebellar sleep spindles.

The analysis of the gait of the mouse during accelerating rotarod learning revealed a progressive adjustment across sessions of the locomotor strategy as evidenced by changes in the combination of gaits used. This proportion varied across trials together with performance improvement, and was then consolidated overnight, indicating long-term retention. The presence of inter-trial variations in motor strategy and variable retention of these changes may reflect the coexistence of two distinct processes, one driving the exploration of different motor strategies and the other allowing for the selective retention of efficient policies. This is consistent with the hypothesis that the discovery of an optimal solution to a given motor learning challenge is driven by an increase in motor variability ^62–64^ followed by the promotion of optimal strategies by reinforcement-learning ^11,16,64,65^.

Reinforcement learning is known to take place at the level of the basal ganglia, notably in the striatum, a structure shown to be critical for accelerating rotarod learning ^30,34^. Here, we further observed that learning progress was associated with a stronger integration of striatal neurons in cerebello-cortical-striatal task-related assembly of neurons, as revealed by canonical correlation analysis. This finding is consistent with the increased role of the DLS in late stages of rotarod learning ^34^ and complements the observation of increased cortico-cerebellar coordination occurring in motor learning ^66^. This stronger integration of striatum in cerebello-cortical-striatal assemblies may be supported by the potentiation of oligosynaptic pathways linking the cerebellum to both M1 and DLS neurons that are evidenced by pairwise correlations and optogenetic stimulations in our study. The sites of plasticity were not identified here, but could involve cerebello-thalamic plasticity ^67,68^ or cortico-striatal plasticity, which has been found to be modulated by the cerebellum ^69,70^.

In this study, we focused on the inter-trial offline activity of the motor network. Indeed, pauses between practice trials has long been known to improve learning^7^, an effect generally associated to consolidation^4,26,27^. Consistently with this, we observed that shortening resting periods in-between trials resulted in a reduced refinement and consolidation of motor strategy, with overall lower performances compared to animals with longer resting periods. During the intertrial periods, we observed both replays of temporal sequences of neuronal activation that occurred in task, as well as reactivations (synchronous activation of task-related ensembles) associated with spindles. Reactivations and replays of temporal sequences of neuronal patterns activation have been extensively associated to the offline consolidation of memory in the hippocampus ^44,47,71^ where replays exhibit strong temporal compression ^72^. Evidence of replays beyond the hippocampus only emerged more recently ^5,43,46,73–76^ but the cerebellum was not examined.

One striking finding is that inter-trial variations of locomotor strategies was in part predicted by the distribution of replays of the neuronal representation of these strategies distributed across the cerebellum, motor cortex, striatum and thalamic nuclei during resting periods between trials. Frequent replays of a given locomotor strategy were found to increase its probability of being used in later trials. Reactivations and replays have been previously linked to planning. Several studies revealed a correlation between hippocampal replays and subsequent trajectories explored in a spatial navigation task ^77^, or with future performances in goal-directed tasks ^78–80^, supporting a potential role in decision making. The effects reported here differ in the sense that the replays do not directly reflect the future strategy but instead correlate with the shift in strategy, and thus more likely reflect a progressive updating process of the future strategy which could help optimize learning. Such offline exploration of the strategy could particularly help the exploration of the possible solutions in complex tasks. Intertrial pauses have indeed been shown to be particularly efficient in learning continuous motor tasks ^81^.

Interestingly, contrary to slow-wave sleep hippocampal or hippocampo-cortical replays, our gait-like replay events were not associated with a clear rhythmic pattern and were not temporally compressed. Such uncompressed replays have already been observed in the forebrain ^43^ but their role had remained elusive. Such recapitulation of activity, with maintained relative timing between neurons, might be a feature of replays involving the cerebellum and could result from the cerebellar contribution to tuning the relative timing of activity on a sub-second timescale ^82^.

Reactivations of task-related hippocampal neuronal activity have also been associated with cortical delta waves ^5,46^ and sleep spindles ^83^. In a previous study, we observed that improvements of performance in the accelerating rotarod are stabilized by an offline consolidation process dependent on the cerebellar output activity ^35^. The contribution of sleep to offline consolidation of skill learning has been indeed widely reported ^84^, and can take place even in short naps taken during the day ^54^. Most notably, sleep spindles occurring during sleep following learning have been associated with improvements in motor performances ^54,56^. Sleep spindles are thought to have a thalamic origin but are known to induce strong cortical activations, either local or widespread. Sleep spindles have been reported only recently in the cerebellum and shown to entrain the motor cortex ^57–59^. We observed that they also entrained neurons in the DLS. Moreover, we found that the retention across sessions of performances and motor strategy correlates with a strong entrainment of task-related cerebello-cortico-striatal assemblies by cerebellar spindle-like events. Moreover, neuronal activity during cerebellar spindles recapitulated the short-term directional interactions between the cerebellum and its target structures observed during the task (see also ^59^). The maintenance across trial of these interactions correlated with the participation of the neurons to cerebellar spindles, consistent with a role of sleep spindles in neuronal plasticity ^28,50,85,86^. Finally, while spindles in the motor cortex have been shown to reactivate task related assemblies ^83^, our results show that such reactivations occur in a much wider network encompassing the cerebellum.

Finally, our work focused on hindlimbs, with neurophysiological recordings located in the corresponding areas of the motor cortex and the dorsolateral striatum, and with an analysis of the kinematic of these limbs. However, the refinement of motor strategy in the accelerating rotarod likely also involve the adjustment of other behavioral parameters such as body postures ^38,39^ which opens further studies on their corresponding neuronal representations and learning mechanisms. Altogether, our results reveal that the exploration and the stabilization of locomotor strategy during motor skill learning rely on distinct but parallel offline neuronal mechanisms which may operate in parallel in time periods between each repetition, and may thus favor learning with “distributed” practice. These mechanisms involve the brain-wide collaboration between different actors of the motor network, which include the cerebellum, the motor cortex and the basal ganglia.

## METHODS

### Animals

Experiments were performed in accordance with the guidelines of the European Community Council Directives. C57BL/6J male mice (6 to 16 weeks old) were kept at a constant room temperature and humidity on 12 hr (non-inverted) light/dark cycle and with ad libitum access to water and food. Behavioral experiments were performed during the light phase.

### Custom accelerating rotarod device

In order to be able to study the behavioral correlates of the accelerating rotarod in mice while performing in-vivo extracellular recordings, we built a custom rotarod device. This rotarod apparatus was made of a single PLA cylinder sharing similar dimensions to the rotarod apparatus commercialized by Bioseb, delimited by two transparent plexiglass walls allowing to contain the mice on the cylinder while being able to observe its limb kinematics during the trial.

During trials, video recording was made through four Imaging Source DMK 37BUX273 grayscale cameras placed around the system to capture the locomotor activity of the mouse in several points of views (left, right, front and back) with an acquisition speed of 50 frames per second. The cylinder was entrained by a brushed DC motor, and the speed of rotation was controlled by an external Arduino micro-controller adjusting the voltage delivered to the motor using a PID control algorithm, reading an optical encoder coupled to the motor to measure instantaneous rotation speed.

The accelerating rotarod protocol was composed of seven consecutive days of training each composed of 7 trials (similarly to the timeline used in ^35^). Rotation speed was initiated at 10 r.p.m. and was incremented by 1 r.p.m. each 8 seconds until reaching 45 r.p.m., with a trial cut-off of 300s. Rotarod trials were followed by a resting period of a minimum of 5 minutes (median time 311s +/- 9 second of median absolute deviation) in which the animal was placed in a plastic enclosure to which it was previously habituated to, in the dark. Each session started and ended by a 20 minute freely moving open-field session.

### Hindpaw tracking and stride classification

Videos acquired during trials were analyzed thanks to a trained neural network on the back view (Deep Lab Cut, Python 3). The position of each hindpaw in the Y axis was considered. The detection of negative peaks in the trajectory of each paw yielded the timing of each paw lift, while the detection of positive peaks yielded the timing of each paw landing. Then, we used the relative timing of swing onsets and offsets of each hind paw to classify each stride in different category. The left hindpaw was taken as a reference. Each swing onset of the left paw was either preceded or followed by the swing onset of the right hindpaw. Bound gaits were events during which the swing onset of the second hindpaw of the stride was preceding the swing offset of the first hindpaw of the stride. However, if the swing onset of the second paw was following the swing offset of the first one, the strides were classified in two other categories. If the closest swing onset of a right paw was preceding the onset of the left hindpaw, the stride was considered as a right gallop type, otherwise it was a left-gallop stride type. Proportion of different strategies was calculated as the number of strides corresponding to a given strategy during a trial divided by the total number of strides during this trial. Within-sessions changes were defined as variations in proportion of a type of gait between the first trial and the last three trials of the session. Across-sessions changes were defined as variation in proportion of a type of gait between the first trial of a session and the first trial of the subsequent session. Consolidation error was calculated as the absolute value of the difference between within-session change in strategy and across-session change in strategy. Because consolidation error is agnostic to the polarity of the initial within-session variation, we also created a consolidation metric, considering the congruence between within-session and across-session changes. It was calculated as the difference between within-session change in strategy and across-session change in strategy, and if the within-session the within-session was negative it was multiplied by −1. The sign of this consolidation metric then reflects the congruence of the of within-session and across-session change, with a value of 0 implying a perfect propagation of the within-session change to the next session, a negative value would reflect an incomplete or incongruent across-session variation, and a positive value would reflect an amplification of the within-session change.

### Electrode design

In order to record neuronal activity in the motor network during the accelerating rotarod learning, we designed a custom made microwire electrode array coupled with an optrode destined to be implanted in the deep cerebellar nuclei (CN). The bundles of electrodes were manufactured by folding in six and twisting a nichrome wire with a 0.005- inch diameter (Kanthal RO-800). Each bundle was then placed inside guide cannulas (10 mm length and 0.16–0.18 mm inner diameter, Coopers Needle Works Limited, UK), glued (Loctite universal glue) to a 3D-printed holder. Individual wires of the bundles were then connected to the electrode interface board (EIB-36; Neuralynx, Boze- man, MT) with one wire for each channel and four channels for each brain region (CL, contralateral VAL, contra- and ipsilateral M1, contra- and ipsilateral DLS), extending 0.5 mm below the tube tip and fixed using gold pins (Neuralynx). In order to perform optogenetic manipulations and recording of the cerebellar nuclei (CN), an optrode was built by associating the bundles destined to be implanted in the CN (eight channels) to an optic fiber (200µm, 0.22NA) housed in a stainless-steel ferrule (Thor labs). Finally, the EIB was secured in place by dental cement. A gold solution (cyanure-free gold solution, Sifco, France) was used for gold plating, and the impedance of each electrode was set to 200–500 kΩ.

### Surgeries

Mice were anesthetized using a mixture of isoflurane and O2 (4% for induction, 1.5% for maintenance). Injections with buprenorphine (0.05 mg/kg, s.c.) were performed to control pain, and core temperature (37°C) was kept with a heating pad. The mice were then fixed in a stereotaxic apparatus (David Kopf Instruments, USA). After a local subcutaneous injection of lidocaine under the scalp (2%, 1 ml), a medial incision was performed, exposing the skull. Small craniotomies were drilled above the recording sites and above the optic fiber location (above the virus injection site). Then, AAV8.hSyn.ChR2(H134R)-mCherry (Addgene catalog#26976-AAV8, 400 nl) was injected into the cerebellar nuclei (CN), followed by a descent of the optrode in the CN, and by the descent of the remaining electrodes in the brain. This procedure allowed us, in one experimental set, to record in the motor cortex bilaterally (M1) (AP 0 mm and –1 mm ML from the Bregma, DV = 0.8 mm depth from the dura), ventrolateral thalamus (VAL) (–1.34 mm AP, ML = 1.00 mm, and DV = 3.6 mm depth from the dura), centrolateral thalamus (CL) (AP at –1.58 mm, ML = 0.9 mm, DV = 3.00 mm depth from the dura), dorsolateral striatum bilaterally (DLS, 0 mm AP,±2.7 mm ML, 2.5 mm depth from dura) and the cerebellar nuclei (CN, 6 mm AP, ±2.3 mm ML, 2.4 mm depth from dura). The ground screw was placed on the surface of the right parietal bone. Super Bond cement (Dental Adhesive Resin Cement, Sun Medical CO, Japan) was applied on the surface of the skull to strengthen the connection between the bone and the cement. We then waited 4 weeks post-surgeries to start the learning protocol, in order to allow for a steady expression of ChR2 in the CN. Placements of electrodes and optic fibers were verified after the end of the protocol.

### Chronic in vivo extracellular recordings

Recordings were performed in awake freely moving mice during the Open field as well as the accelerating rotarod sessions. A custom-made pulley system balanced the weight and torque of the wires during running and allowed the wires to accompany the mouse during the accelerating rotarod task. Signal was acquired using a headstage and amplifier from TDT (32 channels headstage, RZ2, Tucker-Davis Technologies, USA) and analyzed with Python 3.9. Local Field Potential was extracted by downsampling the recorded signal to 1kHz and band-pass filtering it between 0.5Hz and 300Hz (second order bilateral Butterworth filter). The spike sorting was performed using MountainSort4. A total of 5 mice were successfully implanted with the previously mentioned electrode design. Out of these 5 mice, one did not display any functional recording sites in the CN (G024), however, the optic fiber was well positioned in the CN. Thus, this mouse was still considered for the behavioural analysis (Fig. 1 and Fig. 1Sup1) and for the quantification of response to optogenetic stimulations of the CN (Fig. 2Sup1 and Fig. 4). One mouse did not display functional recording sites in M1i and DLSi (G022). Due to technical issues during recording, one learning session (mouse G019) could not be recorded on day 2.

### Optogenetic stimulation of the Cerebellar Nuclei

The injection of an AAV expressing ChR2 coupled with the implantation of an optic fiber in the left CN allowed for the optogenetic activation of CN neurons. Illumination was performed using a LED driver and a 470nm LED (Mightex Systems). Light transmission from the LED output to the implanted optic fiber was done through a 200µm diameter flexible optic fiber. The coupling between the flexible fiber and the implanted optic fiber was allowed by a ceramic mating sleeve. The intensity of current provided to the LED was calibrated in order to obtain an illuminance of 40mW/mm² at the tip of the optic fiber. Optogenetic stimulations of the CN were performed during 100 ms at a frequency of 0.25 Hz during 10 minutes in an Open Field arena, before the start of the rotarod session. The average increase in firing rate during the stimulation was determined by computing the peristimulus time histogram (10 ms bins) of the spikes during the stimulation; the spike count in the histogram was divided by the duration of the bin and the number of stimulations administered to yield a firing rate. The response to stimulation was only analyzed in neurons where at least one bin during the stimulation was 3.5 times larger than the SD of the values of firing rate during baseline (300ms before the onset of stimulation, 10ms bins).

### Putative immobility detection during resting periods

In order to detect bouts of movements during resting periods where no video was recorded, we trained naive Bayesian classifiers to predict the presence of movement based on open recording channels in our electrodes (as they are exquisitely senstitve to movement, and produce large motion artifacts), using the open-field sessions before and after rotarod as ground truth. The ground truth dataset was constituted using the video recordings of the open-fields sessions (30fps videos), processed using the freezing analysis of the EzTrack package (https://github.com/denisecailab/ezTrack) separating bouts of movements and immobility. Then, we processed the recorded signals of open-channels by downsampling it recorded signal to 1kHz and band-pass filtering it between 0.5Hz and 300Hz (simimarly to the Local Field Potential), a Hilbert transform was then applied to extract the instantaneous amplitude of the signal, which was then smoothed using a gaussian kernel of 333ms of standard deviation, yielding a smoothed amplitude reflecting the presence of motion artifacts. Estimations of motion based on the videos were upsampled to 1kHz to match the smoothed amplitudes. Then, mobility and smoothed amplitudes were separated in training and testing datasets in order to cross-validate the classifiers (80% training, 20% testing). We trained Naive Bayesian Classifiers (naive_bayes.GaussianNB from the package scikit-learn) to predict the presence of movements from the smoothed amplitudes. Classifers were trained for each recording sessions of individual mice. In the eventuality where multiple open recording channels would be present on an electrode, a classifier was trained on each of these channels and the classifier with the highest accuracy on the test dataset was selected to detect putative immobility during resting periods. Ultimately, we used the inferred putative immobility during resting to consider only replays occuring outside of movement bouts, and to classify the sleep cycles (see later sections).

### Quantification and statistical analyses

The figures display either mean +/- SEM or boxplot with boxes reporting the lowest and highest quartiles, and whiskers representing 1.5 interquartile range, and points beyond the whiskers are plotted individually as outliers. In general, non-parametric statistical approaches were used for statistical testing in this study, with Mann Whitney U or Kolmogorov-Smirnoff tests comparing unpaired distributions (respectively *mannwhitneyu()* and *ks_2samp()* functions from the Python package *scipy*), Wilcoxon tests to compare paired distributions or difference to zero (*Wilcoxon()* function from the package *scipy*). Corrections of p-values for multiple testing were performed according to the Holm-Sidak method (*multipletests()* function from the package *statsmodels*).

In order to account for the presence of repeated measures per animal, statisticals tests comparing distributions were preceded by a hierarchical nested statistical approach using linear mixed effect models, with random effects corresponding to the different animals fitted as an intercept (*MixedLM()* from the Python package *statsmodels*). The presence of significant dependence between exogenous and endogenous variables was assessed by an ANOVA test on the fitted model (*anova_lm()* function from the package *statsmodels*) followed by a non-parametric test (see Supplementary Tables). This approach was used in Fig. 4c,d,f,g, 5e.

The presence of significant dependence between exogenous and endogenous variables in logistic regressions was assessed by a Chi2 test (*feature_selection.chi2*() function from the package *sklearn*). Linear relationships were assessed using Pearson’s correlation (*pearsonr()* function from the package *scipy*). Difference in proportions between two different conditions were assessed using Fischer exact tests (*fischer_exact()* function from the package *scipy*).

### Detection of replays associated to strides

In order to detect replays of neuronal activity corresponding to a given type of stride during the offline resting period in-between trials, we used a template-matching approach^41–43^. The peri-stimuli time histograms (PSTH, 10ms bins, −1s to 1s centered on left-paw lift) previously computed for each stride type during the sessions were used as templates to perform replay detection during the offline resting period between the trials. The neural activity during resting period for each cell of the network was binned (10ms bins) and centered by subtracting the mean firing rate of each neuron during the resting period, creating a 2-dimensional histogram with similar time scaling as the PSTHs. Then, template matching was perform using the Python package *OpenCV*, which returned a 1-dimensional timeseries of template-matching coefficients. The detection of replays presenting a significant similarity in terms of temporal sequence was done by applying random vertical shuffling on the templates 100 times and performing template matching using these shuffled templates. Vertical shuffling was chosen as a way to detect replays, as it allows for the randomization of the temporal sequence of activity, while preserving co-activation in the network. These shuffled template matchings yielded a mean template-matching coefficient as well as a standard deviation for each timepoint during the resting period. Peaks which were locally higher than 5 SD above the mean were considered as significant replays. As previously described, only replay candidates occurring within periods of putative immobility were considered for the following analysis.

Since patterns of activity may display a degree of resemblance between stride types, candidate replay events matching two stride types were occasionally detected. When replays of different stride type were detected in a time period of 500ms, they were then disambiguated by retaining the replay displaying the highest template-matching coefficient, as it will represent the stride type sharing the highest covariance with the offline activity. To properly quantify our ability to disambiguate template matchings of different gaits in individual replay events, we divided the template-matching coefficient of alternative gaits by the template matching coefficient of the attributed gait. This yielded a disambiguation coefficient, being contained being 0 and 1, with values close to 0 reflecting high ability to disambiguate between the two templates.

In order to assess the similarity between individual gaits and replay events, we implemented the matching index used in ^74^ and in ^75^. For N neurons contained in a recording across brain regions, there would be N(N-1)/2 pairs of neurons. The sequential relationship between their activation was assessed as by comparing their timing of peak firing during individual events (timing of the bin exhibiting the maximum number of spikes). For a given pair of events *a* and *b*, let *m* be the number of neuron pairs with same ordering of peaks firing in events *a* and *b*, and *n* be the number of neurons pairs with dissimilar firing order. The matching index of the pair of events *a* and *b would be defined as follows:*

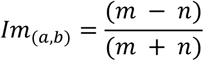

It is a similarity measure bounded by [−1, 1], where a value of 1 reflects an exact same ordering of peak firings in all the pairs of neurons and −1 reflects an exact opposite ordering. This matching index was calculated for every pair of events, either within-gaits (gait x gait), within-gaits (replay x replay) or between gaits and replays (gait x replay).

### Replay-based learning model for motor strategy

Our model assumes that each motor strategy (type of gait) is associated with a “preference” which predicts the probability of being used during the task, and which is updated by the replays of strategy in the intertrial episodes. At each intertrial period, the preference for each gait initially corresponds to the observed proportion of this gait in the previous late trial (second half of the trial). We designed a simple learning model where individual replay events update the gait preference, which can be compared to the actual preference in the next early trial (first half of the trial). The model was elaborated as follows:

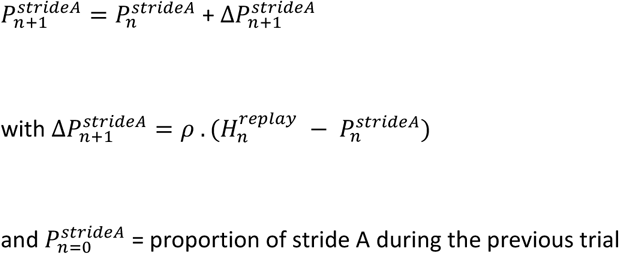

Where 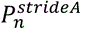 is the predicted preference of using stride A at the replay *n*, *ρ* is the learning rate fitted for each gait/mouse and used for all sessions and intertrials, 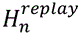 is the updating component, equal to 1 if the replay n was a replay of stride A or equal to 0 otherwise.

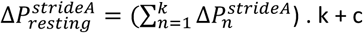

Where 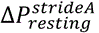 is the variation of proportion of stride predicted by the model during one resting period, and both k and c are other parameters fitted in the model, the former corresponding to a gain and the latter to an offset. This value was then compared to the difference of stride proportion between the early trial following the resting period and the preceding late trial. This model and its three parameters were fitted separately for each mouse and each gait type.

### Canonical correlation analysis

In order to extract common modes of neuronal activity in the motor network during the accelerating rotarod learning, we resorted to the use of canonical correlation, which computes maximally-correlated linear combinations of variables contained in different datasets. Due to our experimental design including simultaneous recordings of the cerebellum, the motor cortex and the dorsolateral striatum, we used the Multiset Canonical Correlation Analysis (MCCA) implemented in the Python package *CCA_zoo*, which allows to apply canonical correlation on more than two datasets simultaneously.

We fitted MCCA on neuronal activity recorded during rotarod trials, with variables being spiketrains of individual neurons binned (10ms) contained within datasets corresponding to their structures (CN, M1 and DLS). The number of components used for the fit of MCCA was determined by the smallest number of neurons recorded in one of the three structures. This fit resulted in the computation of canonical variables for each structure, with each individual neuron being associated to a coefficient for a given MCCA component. These canonical variables exhibited a degree of covariance and represented a fraction of the variance explained from the neuronal activity in their corresponding structure. Ultimately only the first component of the MCCA was considered as it represented the highest fraction of variance explainable by MCCA. Covariance and fraction of explained variance were computed using the *CCA_zoo* package, and these functions were later used for the study of covariance of MCCA neuronal assemblies during resting periods. The global covariance value is obtained from the covariance matrix between the canonical variables of the first component of MCCA, reduced to a single value using singular value decomposition. A high value of covariance thus reflects the presence of a strong common activity of the cerebello-cortico-striatal assemblies.

### Connectivity analysis quantified by sequential activations

Spiketrains of single cells were used to compute cross-correlograms of neuronal pairs. Spiketrains were binned (1ms), then timeseries cross-correlation was applied on the binned spiketrains using the *cross_correlation_histogram()* function from the package *elephant* in order to obtain the cross-correlation of neuronal activity. A central peak around 0 in the crosscorrelogram signals a covariance of firing rate, while an asymmetry signals the presence of sequential activations. The asymmetry of cross-correlogram was quantified using the lag-wise difference in cross-correlation coefficient on a short time window (25ms) preceding and following the spiking of the reference neuron.

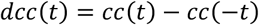

Where *dcc* is the difference in normalized cross-correlation for a given lag value *t*, *cc(t)* is the value of normalized cross-correlation at the lag value *t*, with *t* in the range [1ms, 25ms]. Cumulative asymmetry curved are obtained by computing the cumulative sum of the *dcc(t)*.

Pairs of neurons showing a distribution of lag-wise differences *{dcc}* significantly different from 0 (Wilcoxon rank test, p<0.05) were considered as significantly interacting. If the distribution of differences was significantly higher than 0, the reference neuron was considered as a modulator of the target neuron. A directional coupling index of this asymmetry was computed to reflect the strength of the modulation of the target neuron by the reference neuron, this directional coupling index was calculated as follows:

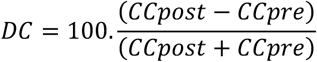

Where DC is the directional coupling index, expressed in percentage, *CCpost* is the average cross-correlogram value in the [1ms, 25ms] time window and *CCpre* is the average cross-correlogram value in the [−25ms, −1ms] time window.

While investigating directional interactions of the cerebellum with motor cortex and striatum, we observed a small number of pairs where an increase in cerebellar firing was followed by a decreased firing in the motor cortex or striatum, resulting in a negative directional coupling index. Due to the small number of these pairs, the analysis in Fig. 4 only focused on pairs with positive directional coupling indices.

### Sleep cycle classification and sleep spindles detection

To identify sleep spindles in the inter-trial rest periods, we first identified the slow-wave sleep as episodes of high delta power in cortical LFP. For this purpose, we split the resting periods into 10s epochs and classified each of these periods using the method described in ^53^. The spectrogram of each 10s epochs of M1 LFP was computed with the Welch method, using the *welch_psd()* function from the package *scipy*. The average power of different frequency bands was then extracted, in the delta band (1-4Hz), in the theta band (5-10Hz) and the average power in the combined bands (2-15Hz). Epochs with high delta power (greater than the average delta power) occuring during putative immobility were considered as NREM sleep, the remaining epochs were considered as wake.

Then, cerebellar sleep spindles were detected within periods of NREM sleep using the following process. The cerebellar recording site displaying the highest average power in the spindle frequency range (10-16Hz) compared to the average power in the wide-band local field potential (1-100Hz) was selected as the site for spindles in the cerebellar nuclei. The signal from this electrode was then filtered in the sleep spindle frequency range (10-16Hz) using a third order bilateral Butterworth filter. A Hilbert’s transform was then applied on this filtered signal, allowing to extract the smoothed amplitude of the envelope of the 10-16Hz signal. The amplitude of the signal during NREM sleep was then z-scored by subtracting its mean and dividing it by its standard deviation. Events where the spindle envelope exceeded 2.5 SD above the mean for at least one sample and the spindle power exceeded 1.5 SD above the mean for at least 300 ms were classified as spindles.

### Sleep spindles locking

The Hilbert’s transform applied on the CN LFP filtered in the range of sleep spindles (10-16Hz) allowed for the extraction of the instantaneous phase of this signal, which was then used to compute the phase of each spike of a given neuron. In order to assess if a neuron was significant phase-locked to cerebellar spindles, we extracted the spikes of this neuron occurring during spindles and their associated phase which allowed for the computation of the Rayleigh Z of the phase distribution using the function *circ_rayleigh()* of the Python package *pingouin*. Then, we iteratively simulated 1000 phase distributions with a number of values equivalent to the number of spikes occurring during spindles, randomly drawn from the instantaneous phase distribution of the signal during spindles, and computed their Rayleigh Z. Neurons displaying a Rayleigh Z higher than the 95^th^ percentile of the distribution of simulated Rayleigh Z were considered as significantly phase-locked to cerebellar spindles.

## Data availability

The dataset generated and analyzed during the current study are available from the corresponding author on reasonable request.

## ACKNOWLEDGMENTS

The authors wish to thank N Alex Cayco Gajic, Valérie Ego-Stengel and Christophe Varin for critical reading of the manuscript, Alexandre Tolboom for his help with Deeplabcut implementation, and Pauline Bonadonna for her help with implant building for multi-sites recording.

## AUTHOR CONTRIBUTIONS

D. P. and C.L. acquired the funding and supervised the project; R.W.S. and A.A. performed electrophysiological recordings. R.W.S performed behavioural experiments, data curation and analysis of electrophysiological and behavioural experiments. All authors interpreted results, participated to the writing and approved the final manuscript.

## FUNDING INFORMATION

This work was supported by Agence Nationale de Recherche to D.P (ANR-19-CE37-0007-01 Multimod, ANR-21-CE37-0025 CerebellEMO, ANR-24-CE37-3944 Ce-Multi-TimeS) and to C.L. (ANR-21-CE16-0036 IntTempComp, ANR-21-CE16-0017 PomPom), by Fondation pour la Recherche Médicale FRM EQU-202103012770 to C.L., by ANR-10-LABX-54 Memolife to R.S. and by the Institut National de la Santé et de la Recherche Médicale (France). This work used the IBENS imaging facility (IMACHEM-IBiSA), member of the French National Research Infrastructure France-BioImaging (ANR-10-INBS-04), which received support from the “Fédération pour la Recherche sur le Cerveau - Rotary International France” (2011) and from the program « Investissements d’Avenir » ANR-10-LABX-54 Memolife.

## CONTACT AND COMPETING INFORMATION

The authors declare that they have no competing interests.

---

**Figure 1Sup1.**
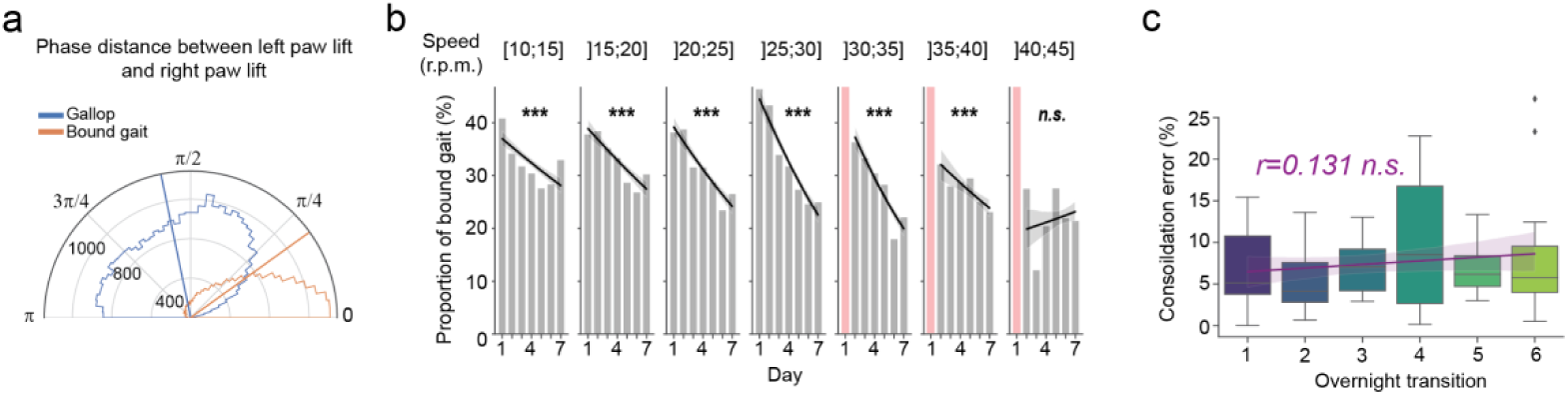
Temporal relationship between paw lifts and time-course of motor strategy adjustment. **a**) Polar histogram displaying the phase distance between the lift of the left hindpaw and the lift of the right hindpaw (relative to the cycle formed by two successive right paw lifts), for bound gait and gallop gaits. Circular means of distributions are represented as vertical lines. **b**) Evolution of the bound gait usage along learning days for each rotarod speed intervals. Lines correspond to logistic regressions. The orange bars signify that the corresponding speed regime was not explored on a particular day ***p<0.001, Chi-square test on logistic regression. **c**) Boxplots displaying the consolidation error of the motor strategy (absolute value of the difference between within-session and across-session variation in motor strategy) along learning days. Linear regression is represented. p>0.05, Pearson’s correlation test. Data from 5 mice, 7 days of accelerating rotarod training with 7 trials/session. See supplementary tables for details on samples and statistics.

**Figure 1Sup2.**
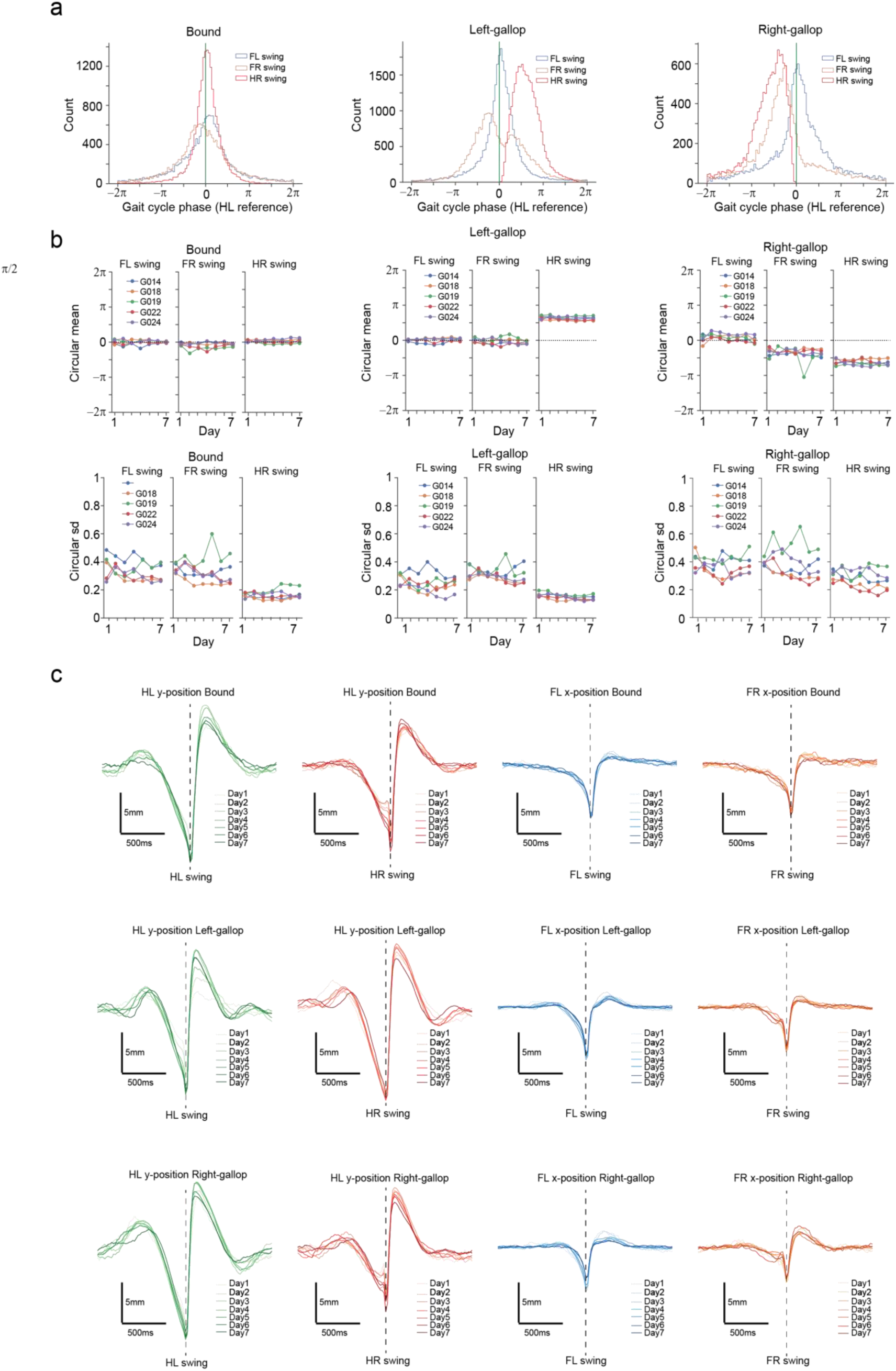
Inter-limb coordination and paw kinematics are stable across learning sessions. **a**) Phase distributions for right hindlimb (HR) and left (FL) and right (FR) forelimb swing onset relative to left hindlimb (HL) swing onset for bound gaits (left), left gallop (middle) and right gallop (right). **b**) Circular means (top) and circular standard deviation (bottom) of limbs swing onset relative to HL paw swing onset for bound gaits (left), left gallop (middle) and right gallop (right). **c**) Median paw kinematics per session centered on the paw swing onset for bound gaits (top), left gallop (center) and right gallop (bottom).

**Figure 1Sup3.**
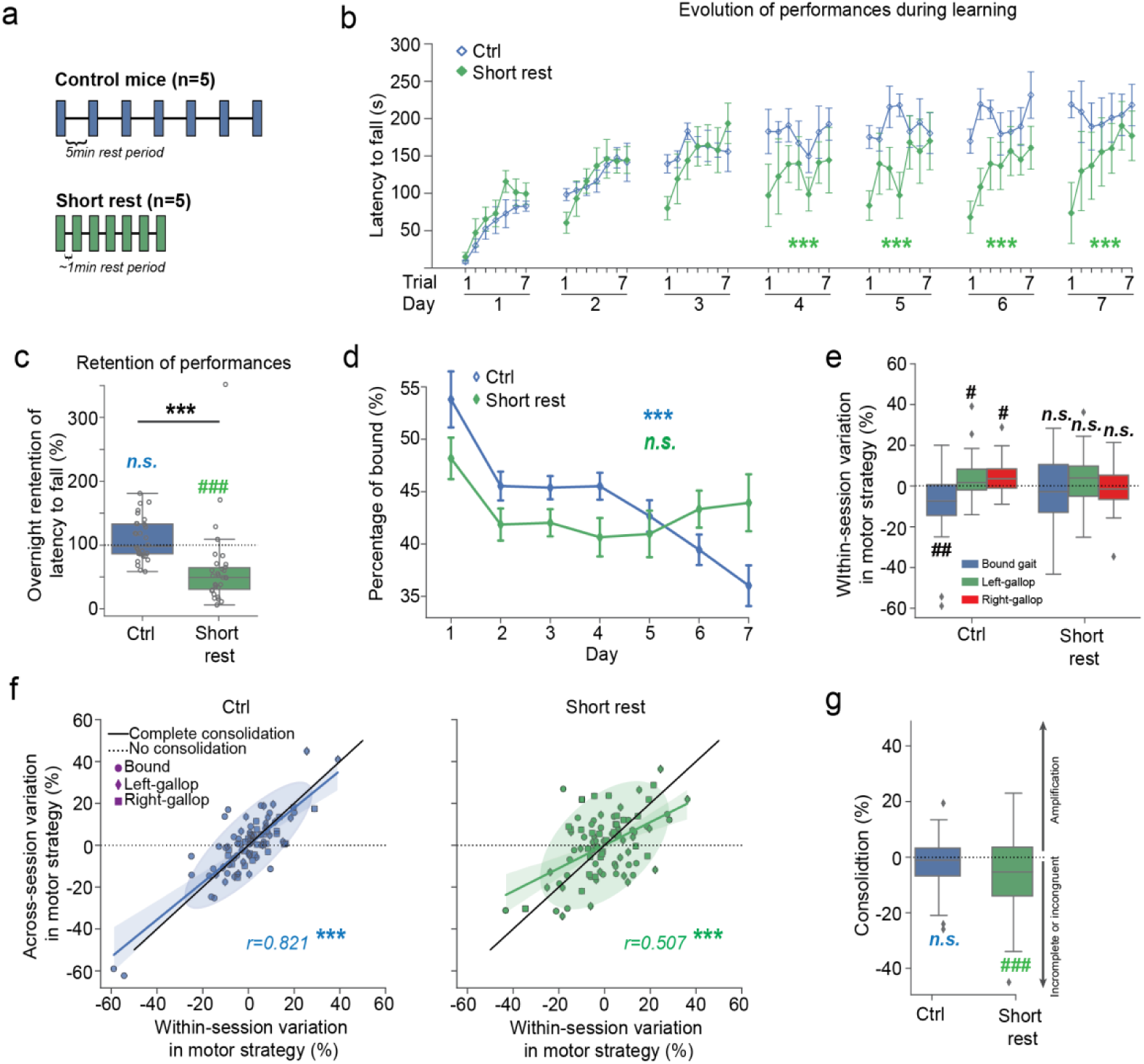
The perturbation of inter-trial resting periods leads to impaired retention of performances, exploration and consolidation of motor strategy. **a**) Experimental design describing a session for control mice (top) and short resting periods (bottom). **b**) Latency to fall during the accelerating rotarod across trials along days for control (n=5) and short rest (n=5) animals, showing a learning deficit of mice undergoing short resting periods. Mixed-effect linear model ANOVA on factor Group, Holm-Sidak corrected. ***p<0.001. **c**) Boxplots displaying the fractional modulation of performances from the last trial of a session to the first trial of a subsequent one, showing an across-session decrease of performances in the short rest group. Mann-Whitney U test, ***p<0.001. Wilcoxon test for difference to 1, ###p<0.001. **d**) Evolution of the percentage of bound locomotion in individual trials along the learning protocol. Mixed-effect linear model ANOVA on factor Day. ***p<0.001. **e**) Boxplots displaying the within-session changes in motor strategy, displaying a systematic decrease of bound gait in control animals but not in animals undergoing short rests. Wilcoxon test for difference to 0 Holm-Sidak corrected, #p<0.05, ###p<0.001. **f**) Scatterplots comparing within-sessions and across-session variations in motor strategy, for control animals (left) and animals undergoing short rests (right), revealing a reduced consolidation of strategy in animals undergoing short rests. Covariance ellipses of 2s.d. and linear regressions are represented. ***p<0.001, Pearson’s correlation test. See supplementary tables for details on samples and statistics.**g**) Boxplots displaying the consolidation of motor strategy for control and short rest animals, displaying an incomplete consolidation for animals undergoing short resting periods. Wilcoxon test for difference to 0 Holm-Sidak corrected, ##p<0.01.

**Figure 2Sup1.**
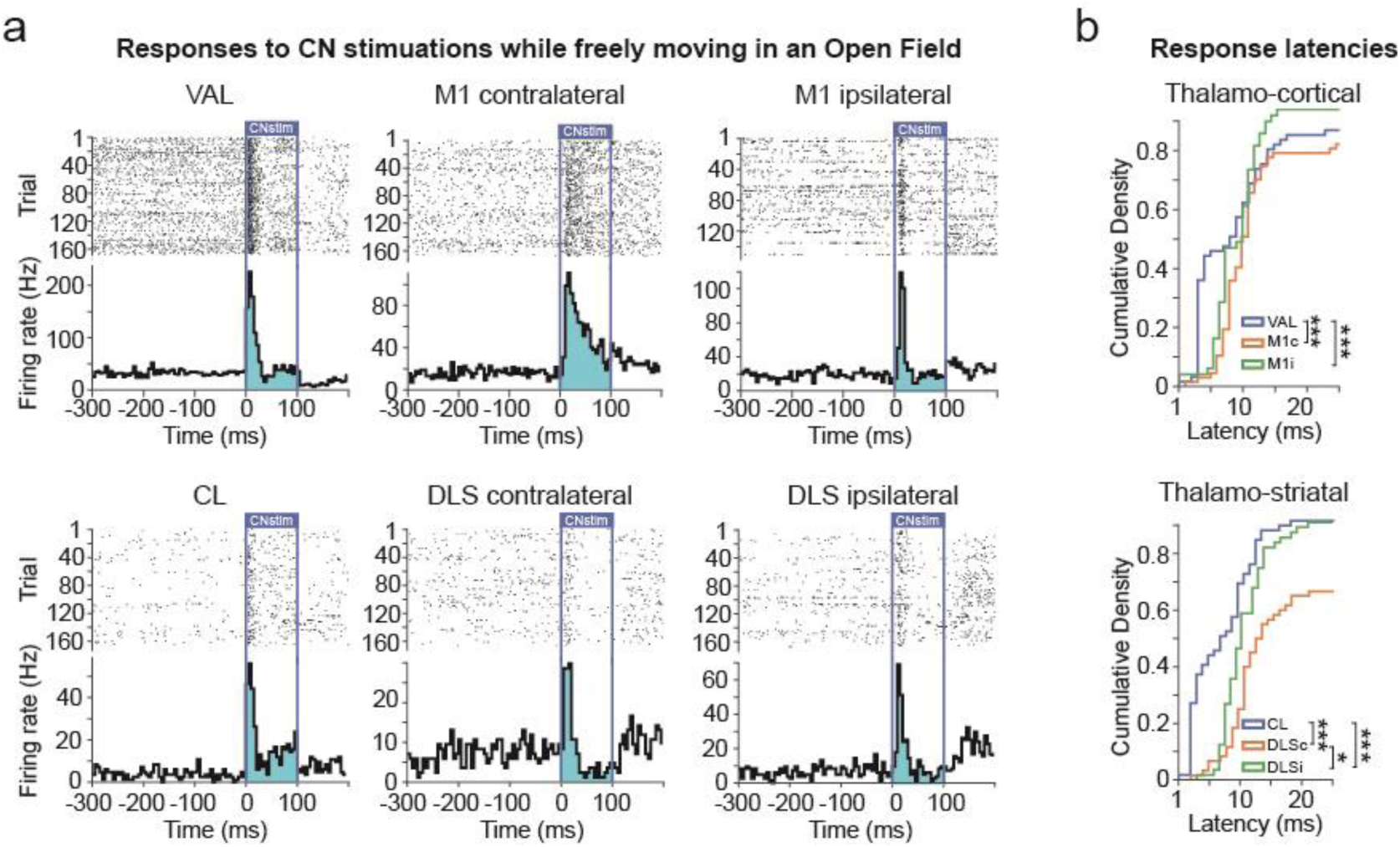
Optogenetic stimulation of the cerebellar nuclei leads to short-latency activation of the thalamus, and of the motor cortex and striatum bilaterally. **a**) Example of peristimulus time histogram (PSTH, 5ms bins) and corresponding raster plots, centered on the onset of the monolateral CN stimulation for single neurons in the VAL, contralateral and ipsilateral M1, CL, contralateral and ipsilateral DLS. **b**) Cumulative histogram displaying the latencies of all responding neurons to optogenetic stimulations in the thalamo-cortical (VAL, contra- and ipsilateral M1: M1c, M1i, top) and thalamo-striatal pathway (CL, contra- and ipsilateral DLS: DLSc, DLSi, bottom). *p<0.05, ***p<0.001, Kolmogorov-Smirnoff test, Holm-Sidak corrected. See supplementary tables for details on samples and statistics.

**Figure 2Sup2.**
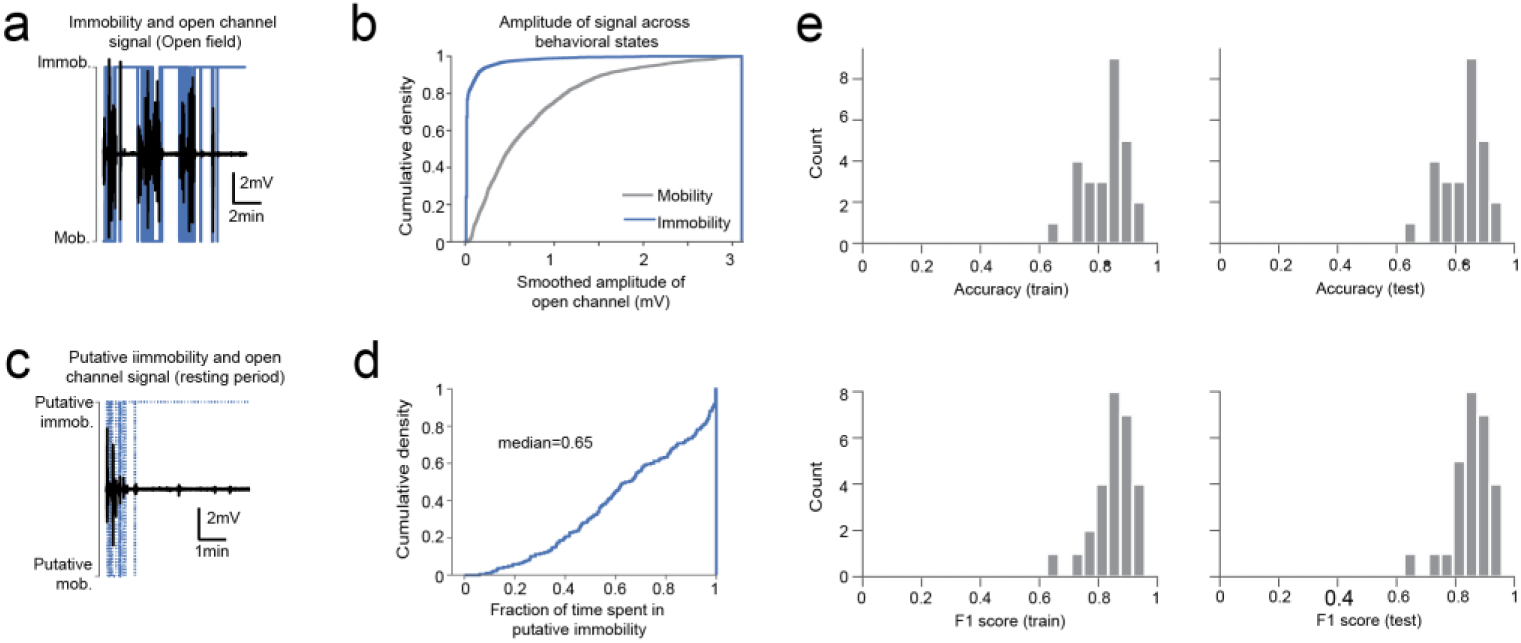
Putative detection of immobility bouts during inter-trial resting periods. **a**) Example of trace recorded on a open channel during an open-field session (black) and video-based detection of mobility (blue), displaying strong movement artifacts during mobility bouts. **b**) Cumulative histogram of smoothed amplitude of recorded signal of the example shown in a), during mobility and immobility. **c**) Example of trace recorded on the same channel as shown in a) during an inter-trial resting period (black) and putative detection of mobility (dotted blue), displaying strong movement artifacts during mobility bouts. **d**) Cumulative histogram of the fraction of time in putative immobility during individual resting periods. **e**) Accuracy (top) and F1 score (bottom) of the cross-validatzed naive Bayesian classifiers trained on open-field session, on training dataset (left, 80% of data points) and test training dataset (right, remaining 20% of data points).

**Figure 2Sup3.**
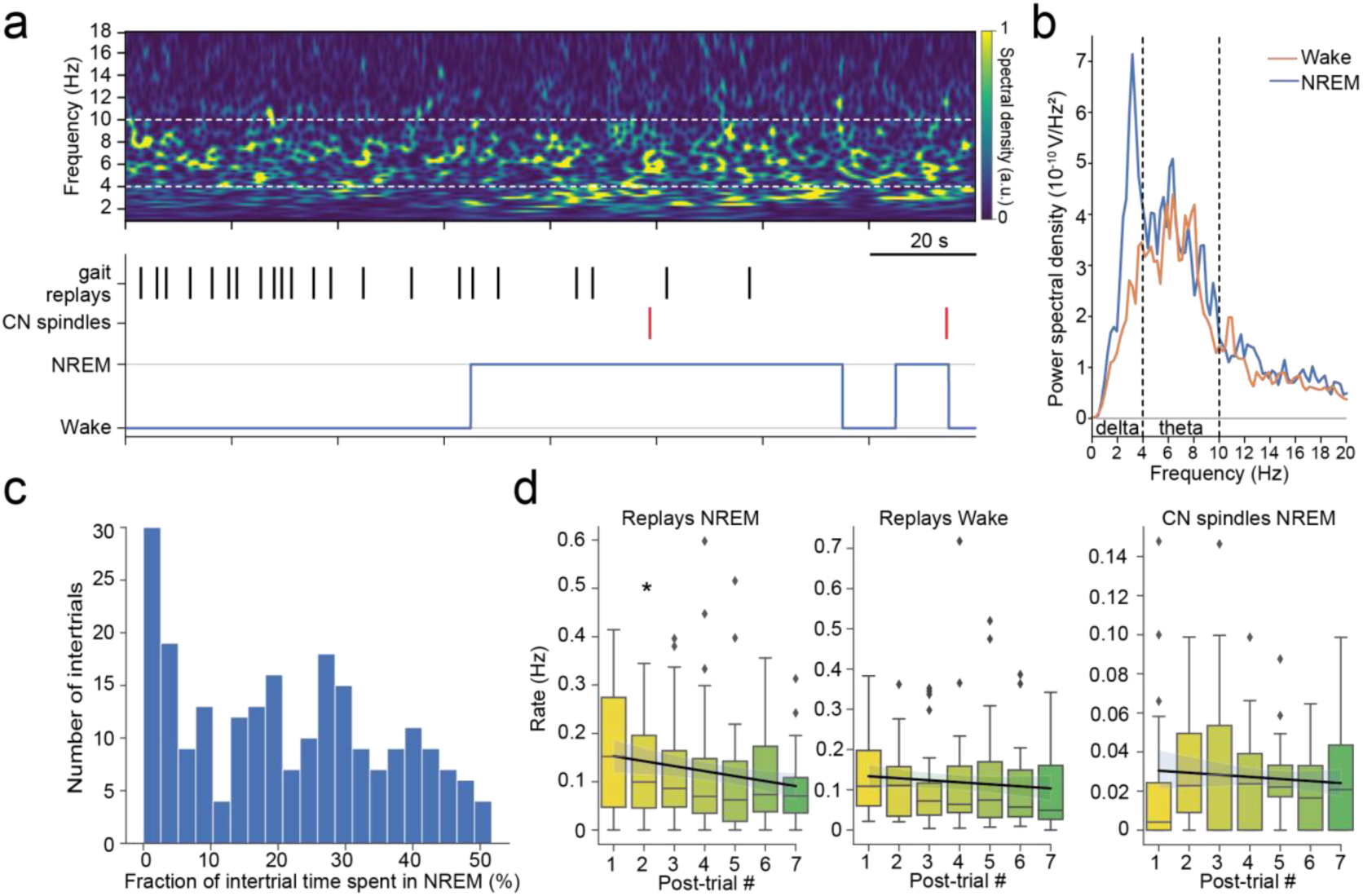
Distribution of replay and spindle events in intertrials. **a**) Example spectrogram of M1 LFP (top) and hypnogram (bottom) in an intertrial period. **b**) Power spectral density of the NREM and and wake intertrial for example a. **c)** Distribution of fraction of time spent in NREM in all intertrial episodes for all mice. **d)** Evolution of rate of replays and sleep spindles in successive intertrials. Pearson’s correlation test: *p<0.05, ***p<0.001. See supplementary tables for details on samples and statistics.

**Figure 2Sup4.**
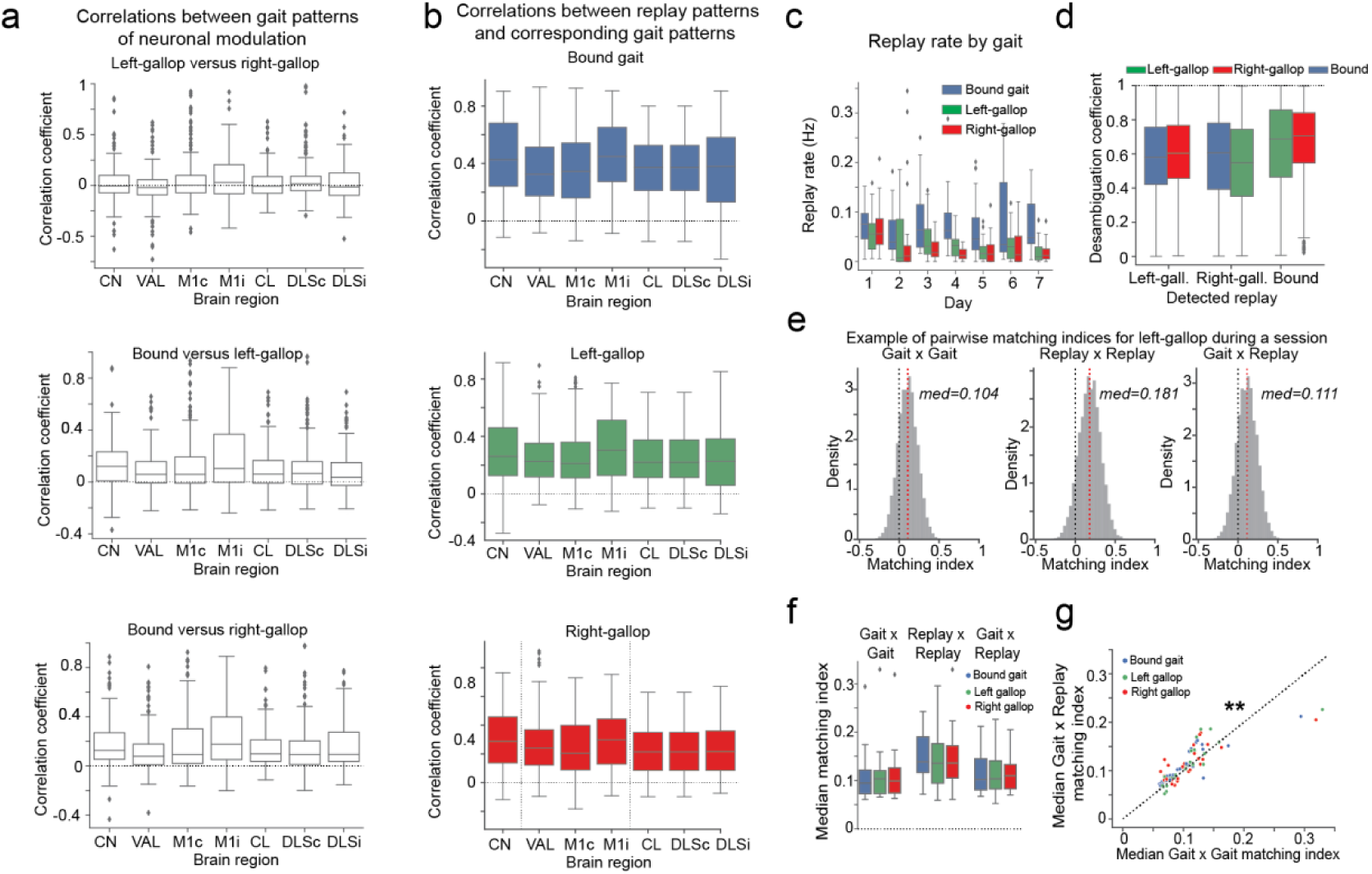
Neurons in the motor network display gait-specific activations during the task, which are faithfully replayed during resting periods. **a)** Correlation coefficients of the average activity of each neuron during different types of gaits. **b)** Correlation coefficients of the average activity of neurons during detected replays compared to their average activity during the corresponding type of gait. **c)** Distribution of replay rates along learning days. Individual data points correspond to single inter-trial rest periods. **d)** Disambiguation coefficients between the detected replays and alternative types of gaits. **e)** Histogram showing the distributions and medians of pairwise matching-indices for left gallop in a training session, separated in within-gait comparisons (left), within-replays comparisons (middle) and gait-replay comparisons (right). **f)** Boxplots showing the distributions of medians matching-indices per sessions for different gaits, separated in within-gait comparisons (left), within-replays comparisons (middle) and gait-replay comparisons (right). **g)** Scatterplot showing the comparison between medians within-gait matching-indices and median gait-replay matching-indices per sessions for different gaits, displaying higher levels of similarity in gait-replay comparisons compared to within-gaits comparisons. See supplementary tables for details on samples and statistics.

**Figure 2Sup5.**
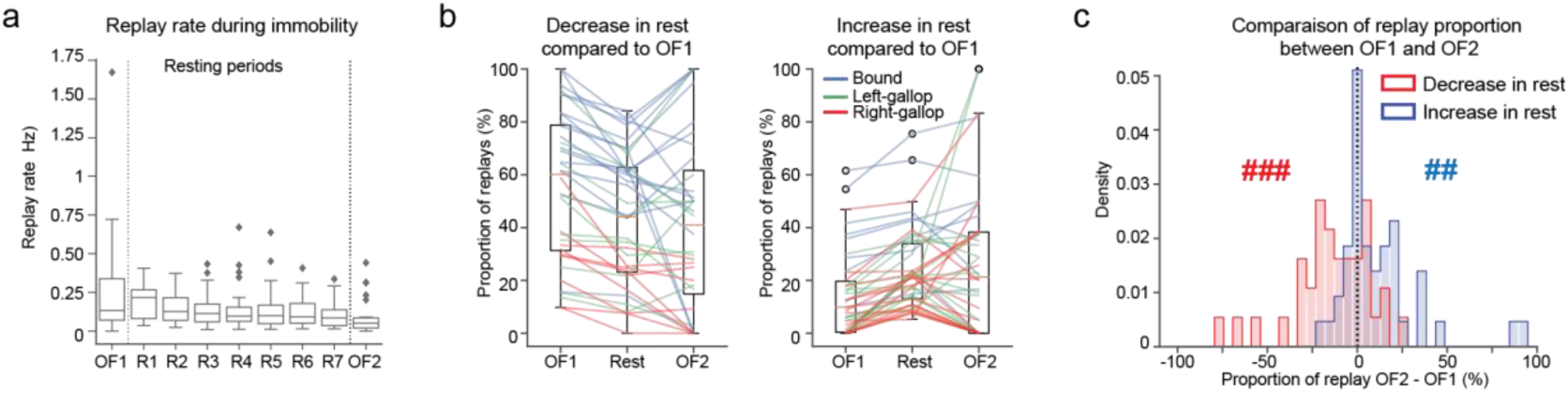
Exposure to the accelerating rotarod paradigm induces long-lasting changes in the structure of gait-specific replays of neuronal activity. **a)** Boxplots displaying the rate of replay detected during immobility episodes in the open-field before (OF1) and after (OF2) the rotarod session, as well as in the episodes of immobility during resting periods in between trials. **b)** Paired boxplots showing the evolution in proportion of different gait specific replays if their proportion decreased during resting period compared to the open-field before the session (left) or if they increased (right). **c)** Histograms showing the difference in proportion of gait specific replays between open field sessions if their proportion decreased during resting period compared to the open-field before the session or if they increased, showing a persistent decrease and increase respectively. ###p<0.001,##p<0.01, Wilcoxon test for the difference to 0. See supplementary tables for details on samples and statistics.

**Figure 2Sup6.**
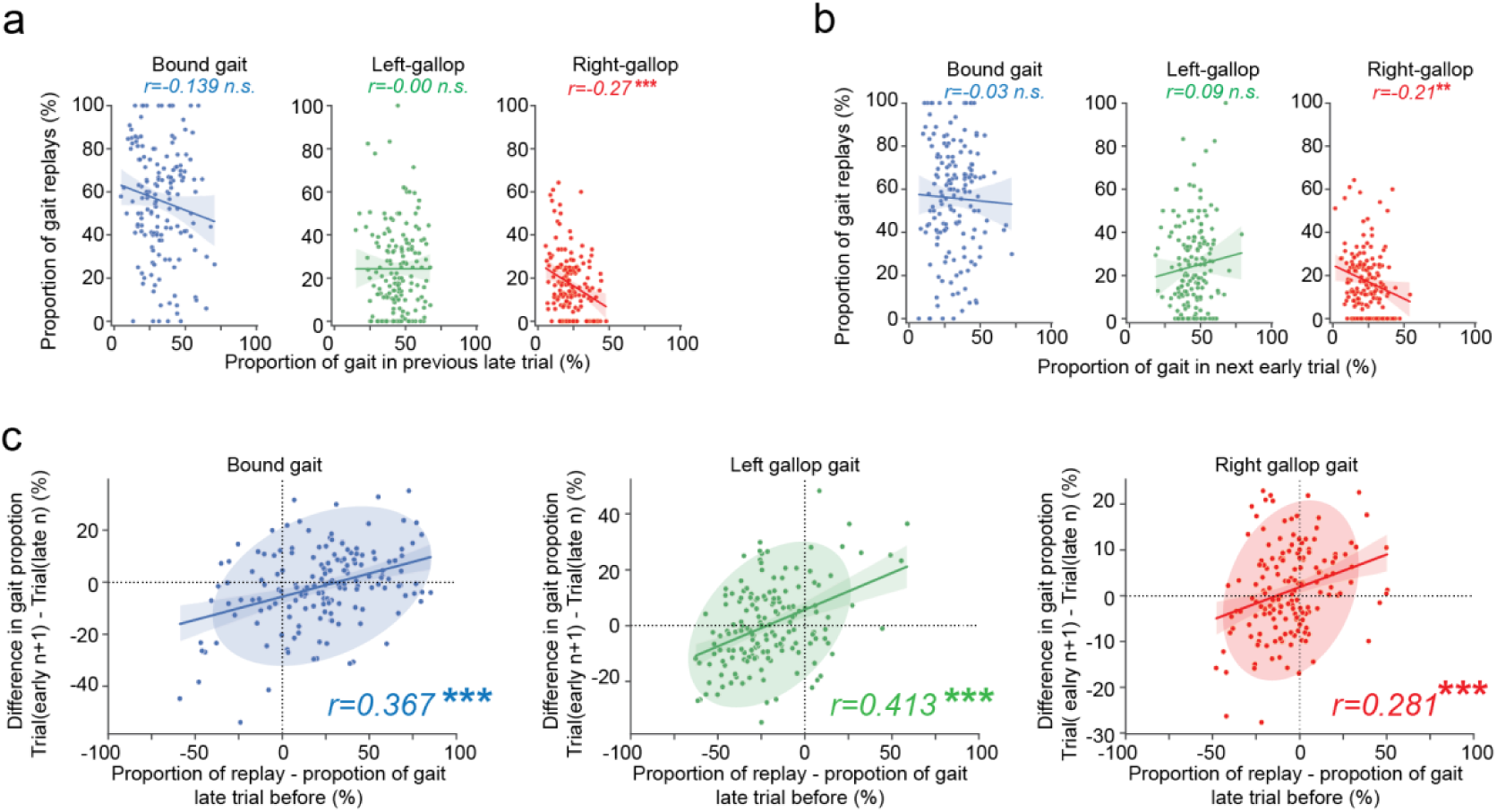
Deviations in proportion of replays compared to the previous usage of a gait correlate with intertrial adjustments of motor strategy. **a)** Scatterplot of the proportion of gait replays and the use of these gaits during the late part of the previous trial. ***p<0.001, Pearson’s correlation test. **b)** Scatterplot of the proportion of different gait replays and the use of these gaits during the early part of the next trial. ***p<0.001, Pearson’s correlation test. **c)** Scatterplot showing the comparison between the deviation in replay proportion from the use of strategy in the late part of the previous trial and the intertrial variation for bound gait, left gallop and right gallop, showing a relationship between the proportion of replays and subsequent strategy adjustments. ***p<0.001, Pearson’s correlation test. See supplementary tables for details on samples and statistics.

**Figure 2Sup7.**
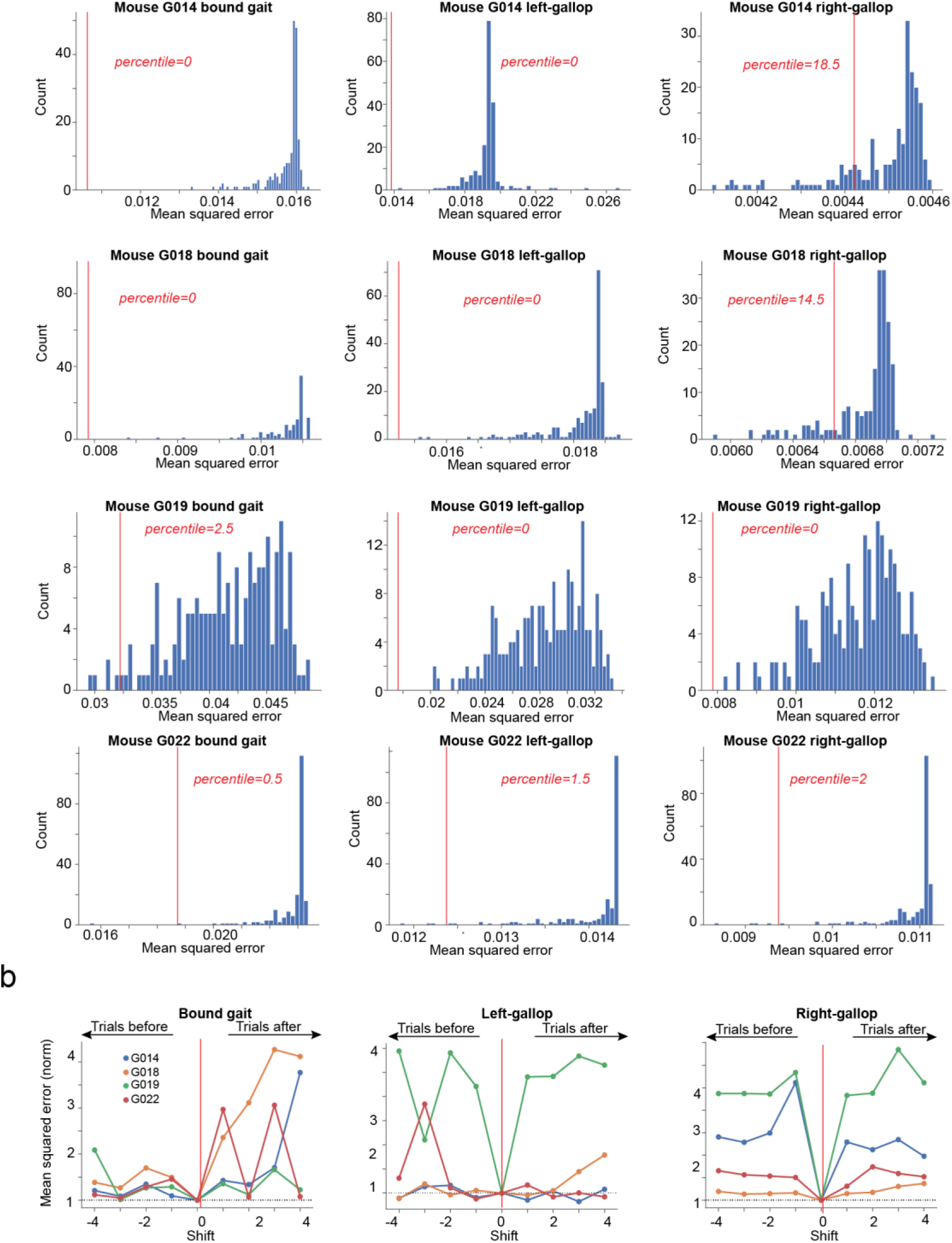
Learning models based on strategy-specific replays trained and evaluated with shuffled, previous or following variations in motor strategy yield less efficient predictions. **a**) Distribution of mean squared errors of models trained with shuffled variations in motor strategy. The vertical red line represents the mean square error of the model trained without shuffling the variations in motor strategy. **b**) Normalized mean squared errors of models trained with previous (negative values) and following (positive values) variations in motor strategy, for bound gait (left), left-gallop (middle) and right-gallop (right).

**Figure 3Sup1.**
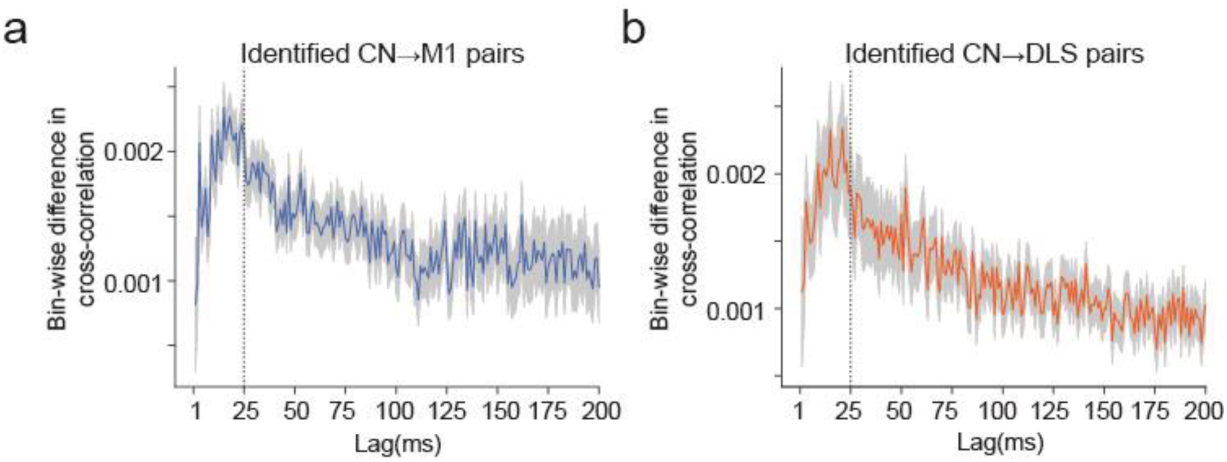
Cerebellar nuclei neurons display short latency directed connectivity with motor cortex and striatum. **a**) Measure of bin-wise asymmetry in normalized cross-correlograms of identified CN->M1 pairs. The vertical dotted line represents the limit of the considered zone for asymmetry detection (25ms). **b**) Same as panel a for CN->DLS pairs. See supplementary tables for details on samples and statistics.

**Figure 3Sup2.**
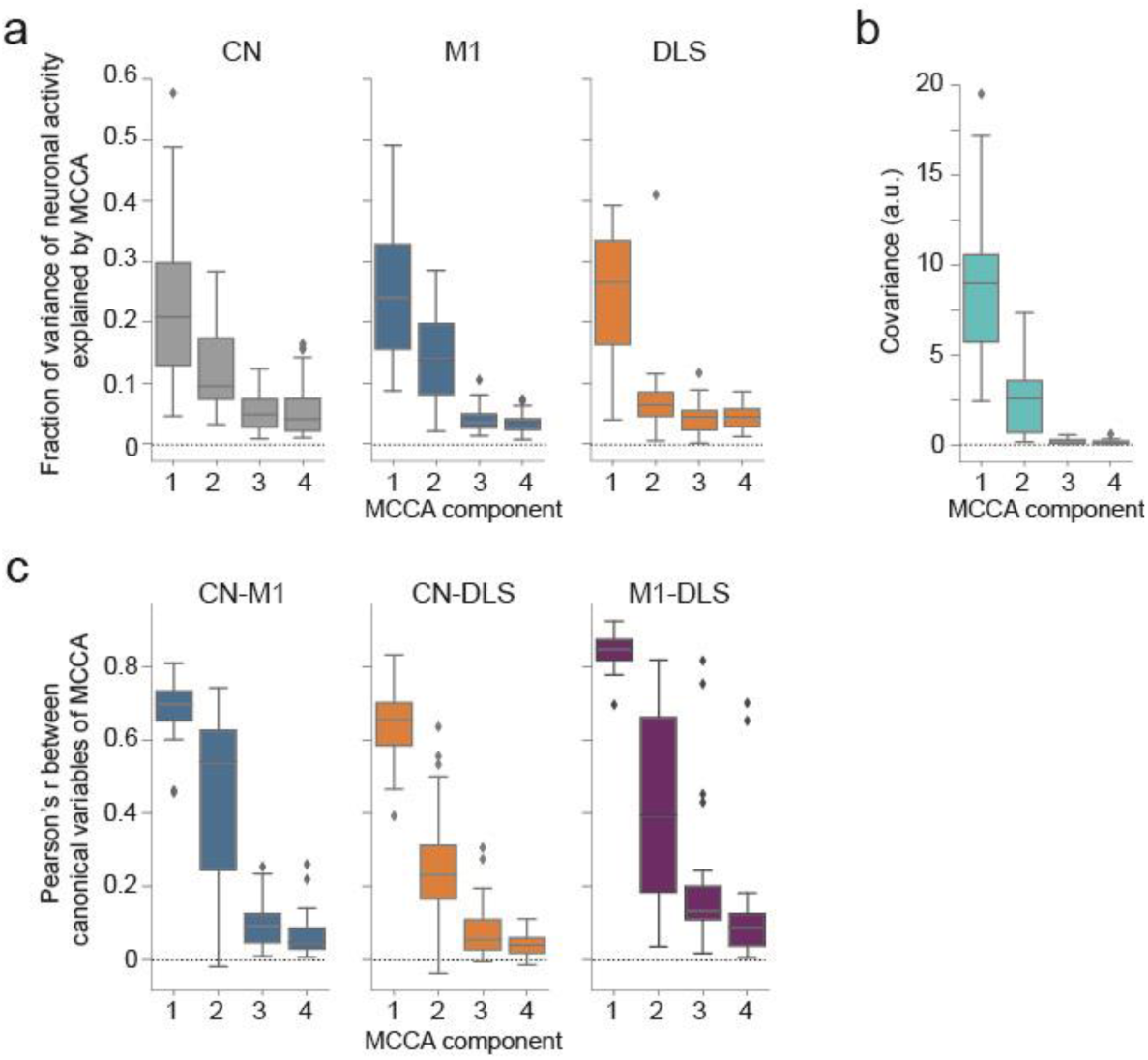
Cerebellum, motor cortex and striatum display common modes of neuronal activity during accelerating rotarod learning. **a**) Distributions of the fraction of variance of neuronal activity captured by MCCA canonical variables in CN (left), M1 (center) and DLS (right). **b**) Distribution of covariance between canonical variables captured by MCCA. **c**) Correlation coefficients between canonical variables captured by MCCA.

**Figure 4Sup1.**
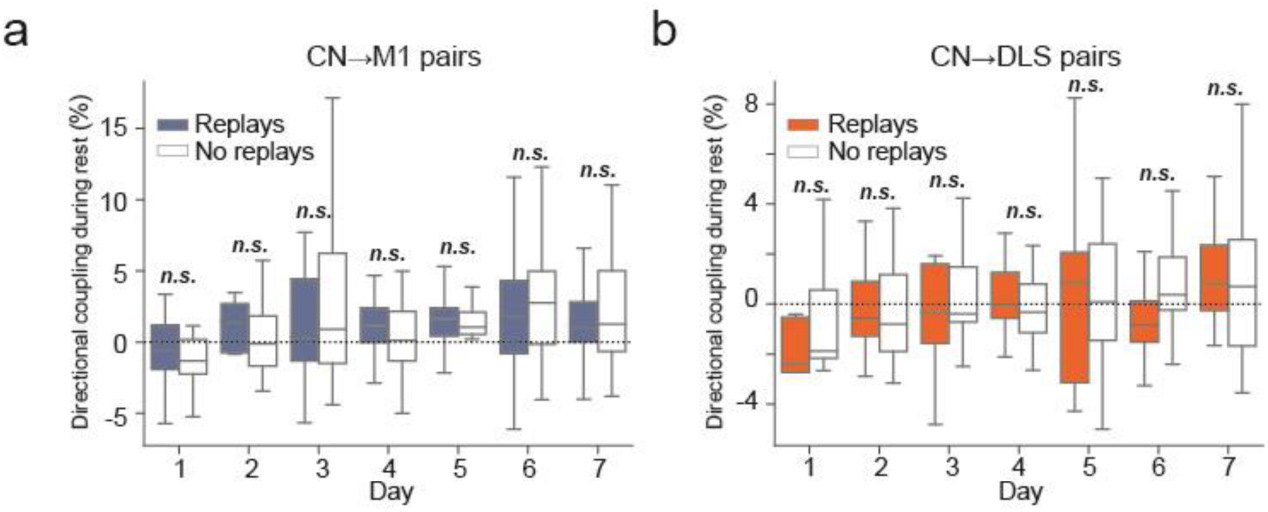
Cerebello-cortical and cerebello-striatal pairs display similar functional connectivity during replays and outside of gait-like replays. **a**) Boxplots displaying the modulation index of CN->M1 pairs during resting periods along learning days, separated between epochs of replays and epochs containing no replays. p>0.05, Wilcoxon test comparing modulation indices during replay to outside of replays. **b**) Same as panel a for CN->DLS pairs. See supplementary tables for details on samples and statistics.

**Figure 4Sup2.**
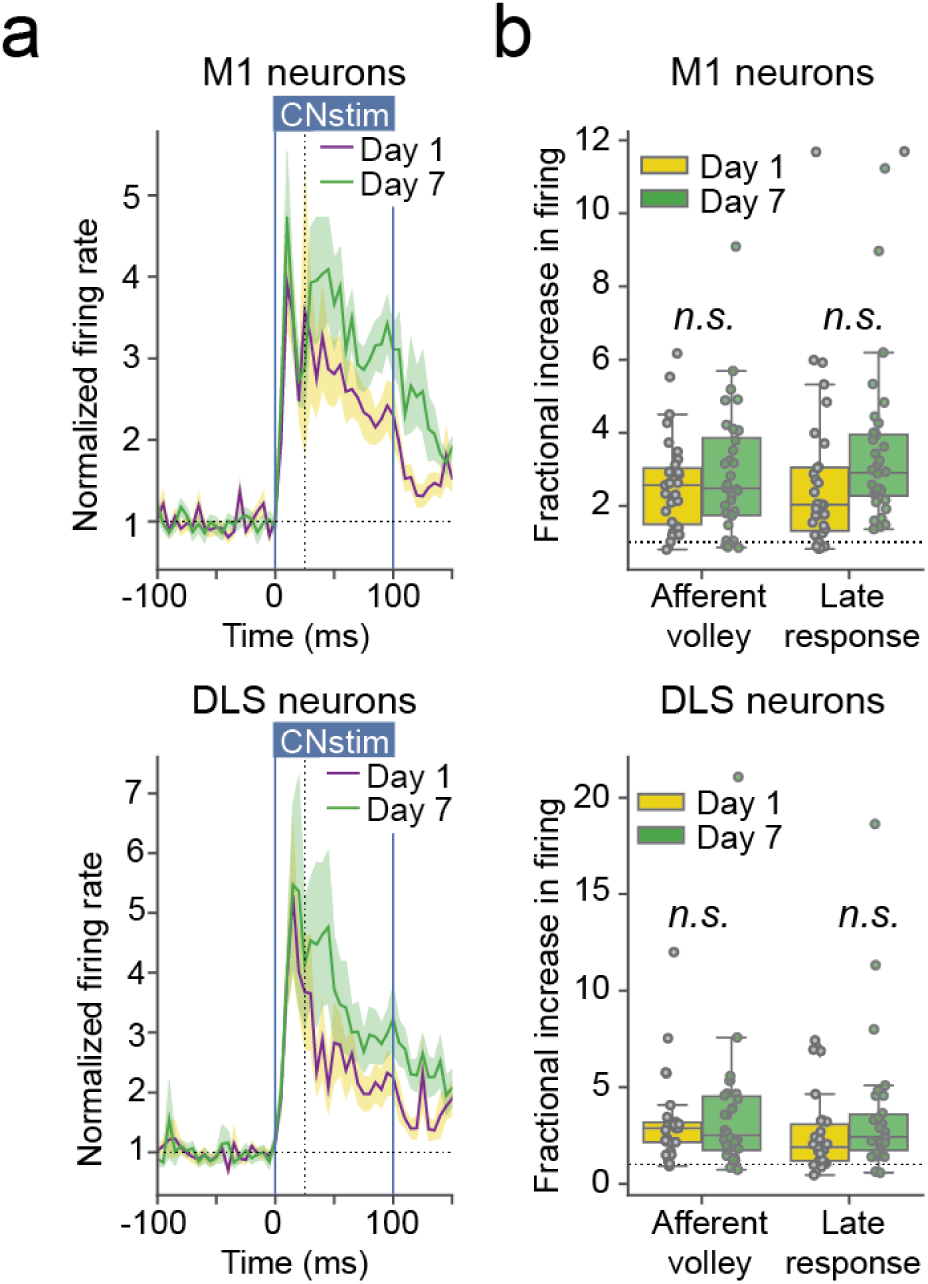
Cerebello-cortical and cerebello-striatal coupling don’t increase in absence of exposure to the accelerating rotarod learning. **a**) PSTH (5ms bins) displaying the change in normalized firing rate (normalized to the 300ms before stimulation onset, average ± SEM) of responsive neurons in M1 (top) and DLS (bottom) in day 1 and day 7 in the absence of exposure to accelerating rotarod learning (vertical dashed lines represent the limit of the afferent volley, 25ms). **b**) Average fractional increase in firing of responsive neurons in M1 (top) and DLS (bottom), during the afferent volley (0 to 25ms) and the late response (25 to 100ms), Mann-Whitney U test. n=4 mice. See supplementary tables for details on samples and statistics.

**Figure 5Sup1.**
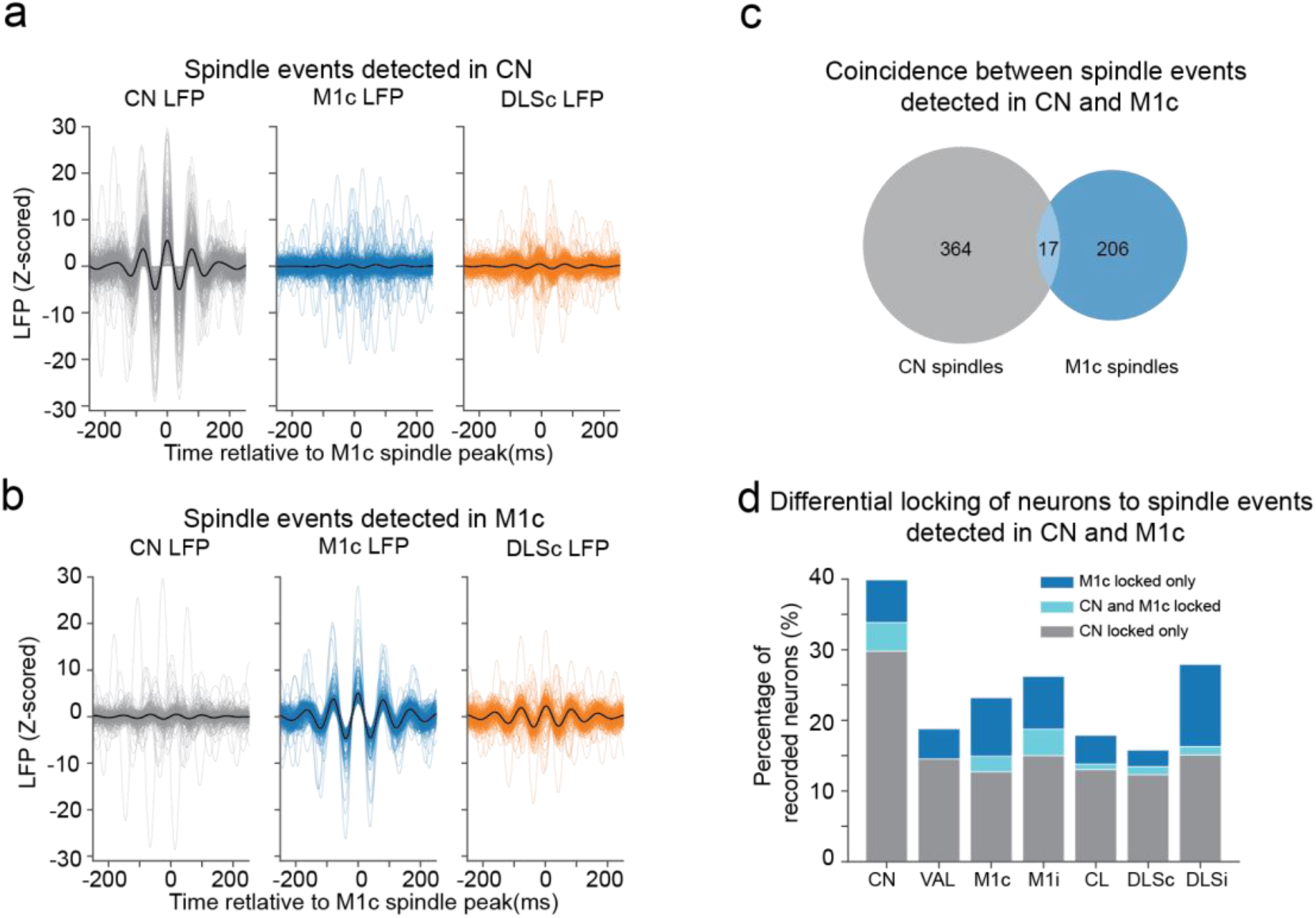
Cerebellar spindles are for the most part distinct from M1c spindles and entrain different neuronal populations. **a**) Overlay of z-scored filtered LFPs (10-16Hz) in the CN (left), contralateral M1 (M1c, centre) and DLS (DLSc, right) centered on CN spindles peak. Black line represents the average LFP. **b**) Same as panel a but aligned on contralateral M1 spindles peak. **c**) Venn diagram displaying rare coincidence between spindles detected in the LFP of CN and M1c. **d**) Proportion of neurons displaying significant locking either to CN spindles only, M1c spindles only, or to both CN and M1c spindles, revealing that CN and M1c spindles tend to engage different populations of neurons.

**Figure 6Sup1.**
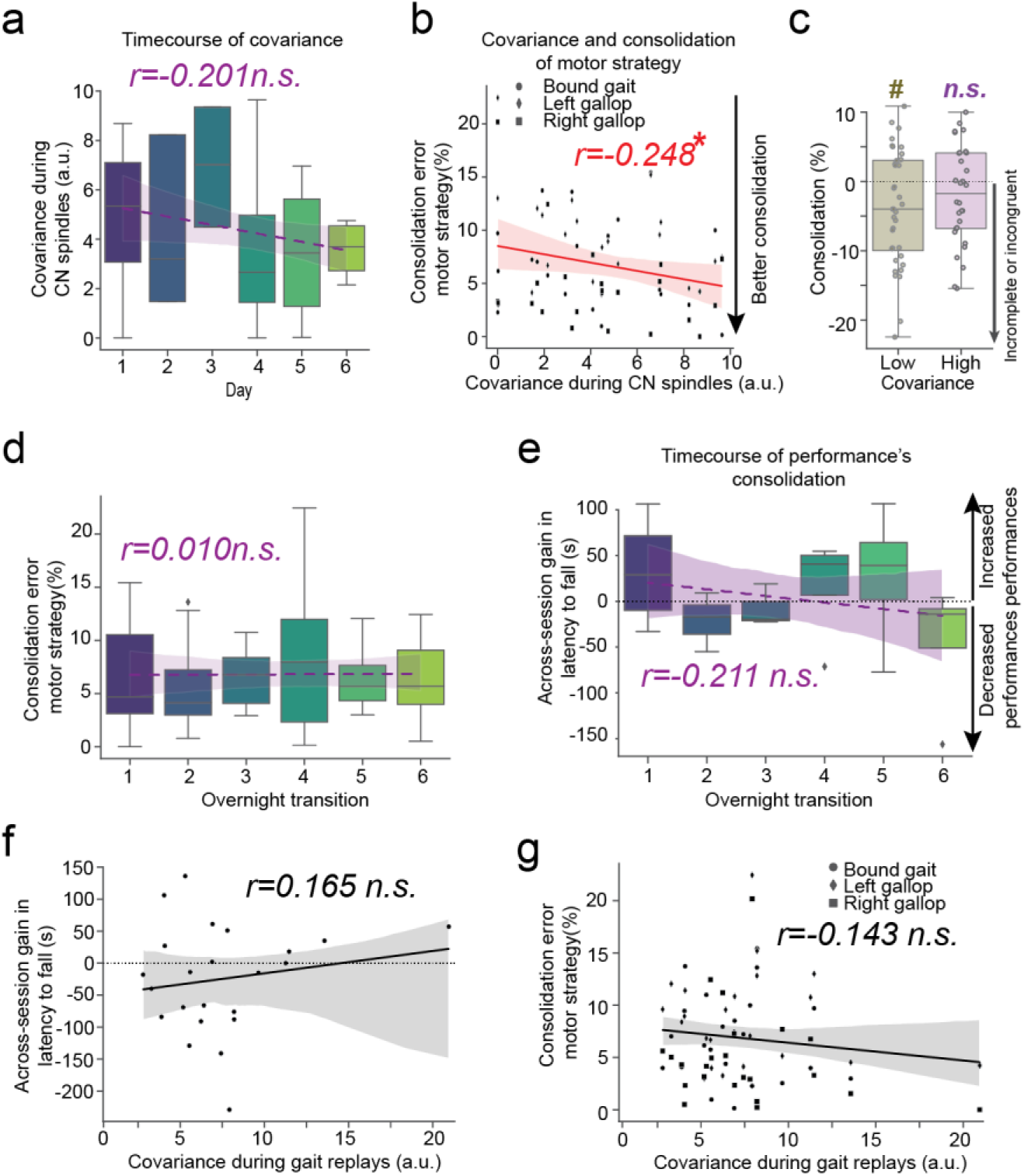
Consolidation of motor strategy and retention of performances don’t display clear time-courses or relationship with covariance within-replays. **a**) Boxplots displaying evolution of covariance during CN spindles along learning days for the mice included in the analysis of figure 6. Linear regression is represented. p>0.05, Pearson’s correlation test. **b**) Scatterplot comparing the error in consolidation of motor strategy to the value of covariance during CN spindles. Linear regression is represented. *p<0.05, Pearson’s correlation test. **c**) Boxplots displaying the consolidation of motor strategy in sessions with low or high covariance during CN spindles. #p<0.05, Wilcoxon test for the difference to 0. **d**) Boxplots displaying the consolidation error of the motor strategy (absolute value of the difference between within-session and across-session change in motor strategy) along learning days for the mice included in the analysis of figure 6. Linear regression is represented. p>0.05, Pearson’s correlation test. **e**) Boxplots displaying the across-session gain in performances (difference between early session performances and late-session performances of the previous session) along learning days for the mice included in the analysis of figure 6. Linear regression is represented. p>0.05, Pearson’s correlation test. **f**) Scatterplot comparing the gain of latency to fall across-session to the value of covariance during gait replays. Linear regression is represented. n.s. not significant, Pearson’s correlation test. **g**) Scatterplot comparing the error in consolidation of motor strategy to the value of covariance during gait replays. Linear regression is represented, n.s. not significant Pearson’s correlation test. See supplementary tables for details on samples and statistics.

**Figure 6Sup2.**
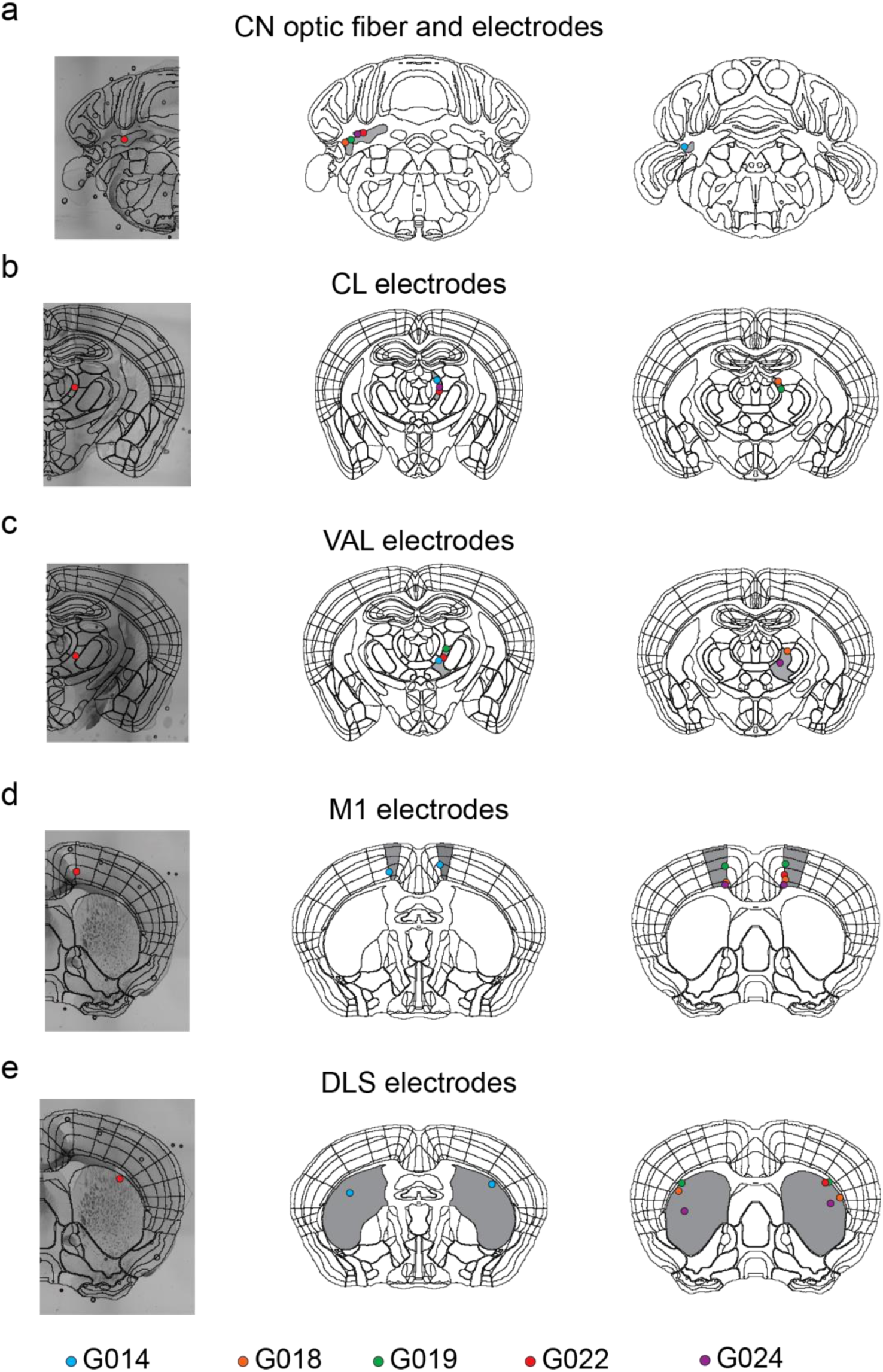
Electrode and optic fiber placements. Example of localization with superimposed delimitation from the Allen atlas (left) and visualization of placement (center and right) for the the optrode implanted in the left CN (**a**), right CL (**b**), right VAL (**c**), left and right hindlimb M1 (**d**) and DLS (**e**). Each color represents individual mouse.

## Notes

### Competing Interest Statement

The authors have declared no competing interest.

