## supplementary tables for "Offline cerebello-cortico-striatal dynamics predict motor strategy exploration and retention in skill learning"

| Day \ n neurons (N mice) | CN | VAL | CL | M1c | M1i | DLSc | DLSi |
|---|---|---|---|---|---|---|---|
| 1 | 23 (4) | 21 (5) | 21 (5) | 30 (5) | 16 (4) | 25 (5) | 16 (4) |
| 2 | 19 (3) | 14 (4) | 17 (4) | 24 (4) | 12 (3) | 25 (4) | 14 (3) |
| 3 | 32 (4) | 21 (5) | 23 (5) | 27 (5) | 16 (4) | 26 (5) | 18 (4) |
| 4 | 24 (4) | 21 (5) | 22 (5) | 25 (5) | 15 (4) | 29 (5) | 20 (4) |
| 5 | 29 (4) | 20 (5) | 21 (5) | 31 (5) | 16 (4) | 26 (5) | 15 (3) |
| 6 | 25 (4) | 23 (5) | 22 (5) | 29 (5) | 16 (4) | 29 (5) | 19 (4) |
| 7 | 28 (4) | 22 (5) | 20 (5) | 26 (5) | 15 (4) | 32 (5) | 16 (4) |

Supplementary Table 1: Statistics for Number of recorded neurons.

| Day \ n pairs (N mice) | CN x M1 | CN x DLS | M1x DLS |
|---|---|---|---|
| 1 | 215 (4) | 203 (4) | 204 (4) |
| 2 | 179 (3) | 191 (3) | 183 (3) |
| 3 | 281 (4) | 280 (4) | 196 (4) |
| 4 | 195 (4) | 236 (4) | 197 (4) |
| 5 | 284 (4) | 219 (4) | 193 (4) |
| 6 | 266 (4) | 261 (4) | 240 (4) |
| 7 | 252 (4) | 310 (4) | 232 (4) |

Supplementary Table 2: Statistics for Number of neuron pairs.

| Day \ n pairs (N mice) | CN->M1 | CN->DLS | M1->DLS |
|---|---|---|---|
| 1 | 35 (4) | 25 (4) | 17 (4) |
| 2 | 22 (3) | 24 (3) | 12 (2) |
| 3 | 54 (4) | 57 (4) | 22 (4) |
| 4 | 34 (4) | 41 (4) | 16 (3) |
| 5 | 38 (4) | 37 (4) | 18 (3) |
| 6 | 89 (4) | 85 (4) | 36 (3) |
| 7 | 40 (4) | 64 (4) | 34 (4) |

Supplementary Table 3: Statistics for Pairs with directional interact.

| N mice | Chi2 | p-value | Sig. |
|---|---|---|---|
| 5 | 511.02 | <0.001 | *** |

Supplementary Table 4: Statistics for Fig1d.

| N mice | Pearson's r | p-value | Sig. |
|---|---|---|---|
| 5 | -0.562 | <0.001 | *** |

Supplementary Table 5: Statistics for Fig1e.

| Time period | N mice | Pearson's r | p-value | Sig. |
|---|---|---|---|---|
| Days 1-3 | 5 | 0.716 | <0.001 | *** |
| Days 4-6 | 5 | 0.402 | 0.122 | n.s. |

Supplementary Table 6: Statistics for Fig1g.

| Model | n (N mice) | Pearson's r | p-value | Sig. |
|---|---|---|---|---|
| Bound | 188 (4) | 0.508 | <0.001 | *** |
| Left gallop | 188 (4) | 0.558 | <0.001 | *** |
| Right gallop | 188 (4) | 0.502 | <0.001 | *** |

Supplementary Table 7: Statistics for Fig2h.

| Category | N mice | Chi2 | p-value | Sig. |
|---|---|---|---|---|
| CN->M1 only | 4 | 7.931 | 0.004 | ** |
| CN->DLS only | 4 | 0.035 | 0.851 | n.s. |
| Both CN->M1 and CN->DLS | 4 | 7220 | 0.007 | ** |

Supplementary Table 8: Statistics for Fig3c.

| Category | N mice | Chi2 | p-value | Sig. |
|---|---|---|---|---|
| CN->M1 | 4 | 4.424 | 0.035 | * |
| CN->DLS | 4 | 4186 | 0.041 | * |

Supplementary Table 9: Statistics for Fig3d.

| Subspace | Pearson's r | p-value | Sig. |
|---|---|---|---|
| CN-M1 | 0.808 | <0.001 | *** |
| CN-DLS | 0.831 | <0.001 | *** |
| M1-DLS | 0.813 | <0.001 | *** |

Supplementary Table 10: Statistics for Fig3e.

| Structure | n (N mice) | Pearson's r | p-value | Sig. |
|---|---|---|---|---|
| CN | 27 (4) | -0.098 | 0.627 | n.s. |
| M1 | 27 (4) | -0.064 | 0.794 | n.s. |
| DLS | 27 (4) | 0.691 | <0.001 | *** |

Supplementary Table 11: Statistics for Fig3f.

| Pair | N mice | Chi2 | p-value | Sig. |
|---|---|---|---|---|
| CN->M1 | 4 | 20176 | <0.001 | *** |
| CN->DLS | 4 | 8788 | 0.003 | ** |

Supplementary Table 12: Statistics for Fig4b.

| Group | Factor | MixedLM ANOVA | p-value | Sig. |
|---|---|---|---|---|
| CN->M1 trials | Day | F(6,160)=2.3826 | 0.0312 | * |
| CN->M1 resting | Day | F(6,160)=4.1132 | <0.001 | *** |
| CN->M1 (trials-resting) | Day | F(6,160)=3.3886 | 0.004 | ** |
| CN->DLS trials | Day | F(6,144)=3.3698 | 0.004 | ** |
| CN->DLS resting | Day | F(6,144)=4.3237 | <0.001 | *** |
| CN->DLS (trials-resting) | Day | F(6,144)=1.1336 | 0.346 | n.s. |

Supplementary Table 13: Statistics for Fig4cd.

| Group | Day | Wilcoxon | p-value (corrected) | Sig |
|---|---|---|---|---|
| CN->M1 resting | 1 | 37 | 0.207 | n.s. |
| | 2 | 1 | <0.001 | ### |
| | 3 | 40 | <0.001 | ### |
| | 4 | 59 | 0.008 | ## |
| | 5 | 18 | <0.001 | ### |
| | 6 | 15 | <0.001 | ### |
| | 7 | 11 | <0.001 | ### |
| CN->M1 (trials-resting) | 1 | 0 | <0.001 | ### |
| | 2 | 14 | 0.033 | # |
| | 3 | 172 | 0.996 | n.s. |
| | 4 | 46 | 0.996 | n.s. |
| | 5 | 132 | 0.887 | n.s. |
| | 6 | 386 | 0.887 | n.s. |
| | 7 | 109 | 0.995 | n.s. |
| CN->DLS resting | 1 | 6 | 0.813 | n.s. |
| | 2 | 43 | 0.378 | n.s. |
| | 3 | 54 | 0.045 | # |
| | 4 | 68 | 0.070 | n.s. |
| | 5 | 55 | 0.102 | n.s. |
| | 6 | 56 | <0.001 | ### |
| | 7 | 60 | 0.002 | ## |
| CN->DLS (trials-resting) | 1 | 0 | 0.137 | n.s. |
| | 2 | 7 | 0.003 | ## |
| | 3 | 28 | 0.002 | ## |
| | 4 | 13 | <0.001 | ### |
| | 5 | 65 | 0.137 | n.s. |
| | 6 | 159 | 0.121 | n.s. |
| | 7 | 126 | 0.137 | n.s. |

Supplementary Table 14: Statistics for Fig4cd(bis).

| Structure | Factor | n (Day 1) | n (Day 7) | Moment | MixedLM ANOVA | p-value | Sig. |
|---|---|---|---|---|---|---|---|
| M1 | Day (1vs7) | 20 | 15 | Afferent volley | F(1,33)=12.2185 | 0.001 | ** |
| | | 20 | 15 | Late response | F(1,33)=1.0015 | 0.324 | n.s. |
| DLS | Day (1vs7) | 13 | 18 | Afferent volley | F(1,29)=7.3061 | 0.011 | * |
| | | 13 | 18 | Late response | F(1,29)=0.9407 | 0.340 | n.s. |

Supplementary Table 15: Statistics for Fig4fg.

| Structure | Moment | Mann Whitney U | p-value | Sig. |
|---|---|---|---|---|
| M1 | Afferent volley | 61 | 0.003 | ** |
| | Late response | 139 | 0.726 | n.s. |
| DLS | Afferent volley | 66 | 0.043 | * |
| | Late response | 95 | 0.389 | n.s. |

Supplementary Table 16: Statistics for Fig4fg(bis).

| Group | Test | n (N mice) | Stat | p-value (corrected) | Sig. |
|---|---|---|---|---|---|
| CN->M1 | Wilcoxon (non-locked) | 95 (4) | 486 | <0.001 | ### |
| | Wilcoxon (locked) | 27 (4) | 110 | 0.059 | n.s. |
| CN->DLS | Wilcoxon (non-locked) | 82 (4) | 425 | <0.001 | ### |
| | Wilcoxon (locked) | 20 (4) | 25 | 0.002 | ## |
| M1->DLS | Wilcoxon (non-locked) | 64 (4) | 147 | <0.001 | ### |
| | Wilcoxon (locked) | 8 (1) | 10 | 0.315 | n.s. |

Supplementary Table 17: Statistics for Fig5d.

| Group | n locked (N mice) | n non-locked (N mice) | Factor | MixedLM ANOVA | p-value | Sig. |
|---|---|---|---|---|---|---|
| CN->M1 | 27 (4) | 95 (4) | Locking | F(1,120)=5.7146 | 0.018 | * |
| CN->DLS | 20 (4) | 82 (4) | Locking | F(1,100)=0.2254 | 0.636 | n.s. |
| M1->DLS | 8 (1) | 64 (4) | Locking | F(1,70)=1.9181 | 0.170 | n.s. |

Supplementary Table 18: Statistics for Fig5d (bis).

| Group | Test | n locked (N mice) | n non-locked (N mice) | Stat | p-value (corrected) | Sig. |
|---|---|---|---|---|---|---|
| CN->M1 | Mann Whitney U | 27 (4) | 95 (4) | 1786 | 0.002 | ** |
| CN->DLS | Mann Whitney U | 20 (4) | 82 (4) | 768 | 0.664 | n.s. |
| M1->DLS | Mann Whitney U | 8 (1) | 64 (4) | 316 | 0.293 | n.s. |

Supplementary Table 19: Statistics for Fig5d (ter).

| n (N mice) | Wilcoxon | p-value (corrected) | Sig |
|---|---|---|---|
| 22(4) | 96 | 0.0434 | # |

Supplementary Table 20: Statistics for Fig6b.

| n (N mice) | Pearson's r | p-value | Sig. |
|---|---|---|---|
| 22 (4) | 0.450 | 0.0361 | * |

Supplementary Table 21: Statistics for Fig6c.

| Category | n (N mice) | Pearson's r | p-value | Sig. |
|---|---|---|---|---|
| Low covariance | 36 (4) | 0.576 | <0.001 | *** |
| High covariance | 30 (4) | 0.789 | <0.001 | *** |

Supplementary Table 22: Statistics for Fig6d.

| Speed bin | N mice | Chi2 | p-value (corrected) | Sig. |
|---|---|---|---|---|
| [10;15] | 5 | 61375 | <0.001 | *** |
| ]15;20] | 5 | 63369 | <0.001 | *** |
| ]20;25] | 5 | 92011 | <0.001 | *** |
| ]25;30] | 5 | 124942 | <0.001 | *** |
| ]30;35] | 5 | 61157 | <0.001 | *** |
| ]35;40] | 5 | 6268 | 0.024 | * |
| ]40;45] | 5 | 0.420 | 0.517 | n.s. |

Supplementary Table 23: Statistics for Fig1Sup1b.

| n (N mice) | Pearson's r | p-value | Sig. |
|---|---|---|---|
| 30 (5) | 0.131 | 0.215 | n.s. |

Supplementary Table 24: Statistics for Fig1Sup1c.

| Day | Factor | MixedLM ANOVA | p-value (Holm-SIdak corrected) | Sig. |
|---|---|---|---|---|
| 1 | Group | F(1,68)=4.1848 | 0.128 | n.s. |
| 2 | Group | F(1,68)=0.0618 | 0.804 | n.s. |
| 3 | Group | F(1,68)=1.7259 | 0.349 | n.s. |
| 4 | Group | F(1,68)=34.9797 | <0.001 | *** |
| 5 | Group | F(1,68)=31.4809 | <0.001 | *** |
| 6 | Group | F(1,68)=49.2714 | <0.001 | *** |
| 7 | Group | F(1,68)=25.4609 | <0.001 | *** |

Supplementary Table 25: Statistics for Fig1Sup3b.

| Group | Test | n (N mice) | Stat | p-value | Sig. |
|---|---|---|---|---|---|
| Ctrl | Wilcoxon | 30 (5) | 158 | 0.198 | n.s. |
| Short rest | Wilcoxon | 30 (5) | 28 | <0.001 | ### |

Supplementary Table 26: Statistics for Fig1Sup3c.

| Test | Group1 | Group2 | n1 (N mice) | n2 (N mice) | Stat | p-value | Sig. |
|---|---|---|---|---|---|---|---|
| Mann Whitney U | Ctrl | Short rest | 30 (5) | 30 (5) | 808 | <0.001 | *** |

Supplementary Table 27: Statistics for Fig1Sup3c(bis).

| Group | Factor | MixedLM ANOVA | p-value | Sig. |
|---|---|---|---|---|
| Ctrl | Day | F(1,243)=61.9398 | <0.001 | *** |
| Short rest | Day | F(1,243)=1.3019 | 0.255 | n.s. |

Supplementary Table 28: Statistics for Fig1Sup3d.

| Group | Pattern | Test | n (N mice) | Stat | p-value | Sig. |
|---|---|---|---|---|---|---|
| Ctrl | Bound | Wilcoxon | 30 (5) | 92 | 0.009 | ## |
| | Left-gallop | Wilcoxon | 30 (5) | 133 | 0.040 | # |
| | Right-gallop | Wilcoxon | 30 (5) | 108 | 0.017 | # |
| Short rest | Bound | Wilcoxon | 30 (5) | 203 | 0.556 | n.s. |
| | Left-gallop | Wilcoxon | 30 (5) | 188 | 0.371 | n.s. |
| | Right-gallop | Wilcoxon | 30 (5) | 216 | 0.746 | n.s. |

Supplementary Table 29: Statistics for Fig1Sup3e.

| Group | n (N mice) | Pearson's r | p-value | Sig. |
|---|---|---|---|---|
| Ctrl | 90 (5) | 0.821 | <0.001 | *** |
| Short rest | 90 (5) | 0.507 | <0.001 | *** |

Supplementary Table 30: Statistics for Fig1Sup3f.

| Group | Test | n (N mice) | Stat | p-value | Sig. |
|---|---|---|---|---|---|
| Ctrl | Wilcoxon | 90 (5) | 1539 | 0.082 | n.s. |
| Short rest | Wilcoxon | 90 (5) | 1180 | <0.001 | ### |

Supplementary Table 31: Statistics for Fig1Sup3g.

| Structure 1 | Structure 2 | n1 neurons (N mice) | n2 neurons (N mice) | KS stat | p-value | Sig |
|---|---|---|---|---|---|---|
| VAL | M1c | 61 (5) | 67 (5) | 0.414 | <0.001 | *** |
| VAL | M1i | 61 (5) | 49 (4) | 0.401 | <0.001 | *** |
| M1c | M1i | 67 (5) | 49 (4) | 0.147 | 0.050 | n.s. |
| CL | DLSc | 59 (5) | 60 (5) | 0.444 | <0.001 | *** |
| CL | DLSi | 59 (5) | 56 (4) | 0.422 | <0.001 | *** |
| DLSc | DLSi | 60 (5) | 56 (4) | 0.271 | 0.022 | * |

Supplementary Table 32: Statistics for Fig2Sup1b.

| Event | n (N mice) | Pearson's r | p-value | Sig |
|---|---|---|---|---|
| Replays NREM | 159 (4) | -0.172 | 0.030 | * |
| Replays Wake | 159 (4) | -0.084 | 0.295 | n.s. |
| CN spindles NREM | 159 (4) | -0.071 | 0.372 | n.s. |

Supplementary Table 33: Statistics for Fig2Sup3d.

| Group1 | Group2 | Test | n (N mice) | Stat | p-value | Sig. |
|---|---|---|---|---|---|---|
| Gait x Gait | Gait x Replay | Wilcoxon | 80 (4) | 784 | 0.002 | ** |

Supplementary Table 34: Statistics for Fig2Sup4g.

| Group | Test | n (N mice) | Stat | p-value | Sig. |
|---|---|---|---|---|---|
| Decrease | Wilcoxon | 36 (4) | 107 | <0.001 | *** |
| Increase | Wilcoxon | 42 (4) | 117 | 0.001 | ** |

Supplementary Table 35: Statistics for Fig2Sup5c.

| Pattern | n (N mice) | Pearson's r | p-value | Sig. |
|---|---|---|---|---|
| Bound | 188 (4) | -0.139 | 0.081 | n.s. |
| Left gallop | 188 (4) | -0.000 | 0.998 | n.s. |
| Right gallop | 188 (4) | -0.271 | <0.001 | *** |

Supplementary Table 36: Statistics for Fig2Sup6a.

| Pattern | n (N mice) | Pearson's r | p-value | Sig. |
|---|---|---|---|---|
| Bound | 188 (4) | -0.033 | 0.081 | n.s. |
| Left gallop | 188 (4) | 0.092 | 0.998 | n.s. |
| Right gallop | 188 (4) | -0.207 | 0.009 | ** |

Supplementary Table 37: Statistics for Fig2Sup6b.

| Pattern | n (N mice) | Pearson's r | p-value | Sig. |
|---|---|---|---|---|
| Bound | 188 (4) | 0.367 | <0.001 | *** |
| Left gallop | 188 (4) | 0.413 | <0.001 | *** |
| Right gallop | 188 (4) | 0.281 | <0.001 | *** |

Supplementary Table 38: Statistics for Fig2Sup6c.

| Pair | Day | Wilcoxon | p-value (corrected) | Sig. |
|---|---|---|---|---|
| CN->M1 | 1 | 55 | 0.874 | n.s. |
| | 2 | 38 | 0.838 | n.s. |
| | 3 | 148 | 0.874 | n.s. |
| | 4 | 128 | 0.869 | n.s. |
| | 5 | 130 | 0.874 | n.s. |
| | 6 | 367 | 0.869 | n.s. |
| | 7 | 84 | 0.874 | n.s. |
| CN->DLS | 1 | 3 | 0.846 | n.s. |
| | 2 | 63 | 0.994 | n.s. |
| | 3 | 136 | 0.994 | n.s. |
| | 4 | 102 | 0.746 | n.s. |
| | 5 | 101 | 0.981 | n.s. |
| | 6 | 123 | 0.746 | n.s. |
| | 7 | 124 | 0.994 | n.s. |

Supplementary Table 39: Statistics for Fig4Sup1ab.

| Pair | Test | Wilcoxon | p-value | Sig. |
|---|---|---|---|---|
| CN->M1 sessions pooled | Directional coupling | 4471 | <0.001 | *** |
| CN->DLS sessions pooled | Directional coupling | 4351 | 0.345 | n.s. |

Supplementary Table 40: Statistics for Fig4Sup1ab(bis).

| Structure | Factor | n (Day 1) | n (Day 7) | Moment | MixedLM ANOVA | p-value | Sig. |
|---|---|---|---|---|---|---|---|
| M1 | Day (1vs7) | 31 | 32 | Afferent volley | F(1,61)=1.2367 | 0.270 | n.s. |
| | | 31 | 32 | Late response | F(1,61)=2.4736 | 0.121 | n.s. |
| DLS | Day (1vs7) | 26 | 30 | Afferent volley | F(1,54)=0.2272 | 0.636 | n.s. |
| | | 26 | 30 | Late response | F(1,54)=1.7585 | 0.190 | n.s. |

Supplementary Table 41: Statistics for Fig4Sup2b.

| Structure | p(CN-locked) x p(M1-locked) | n CN/M1-locked | n total | Binomial test | p-value | Sig. |
|---|---|---|---|---|---|---|
| CN | 0.034 | 8 | 198 | 0.040 | 0.557 | n.s. |
| VAL | 0.006 | 0 | 117 | 0.000 | 1 | n.s. |
| M1c | 0.016 | 4 | 181 | 0.022 | 0.373 | n.s. |
| M1i | 0.022 | 3 | 80 | 0.375 | 0.239 | n.s. |
| CL | 0.007 | 1 | 123 | 0.008 | 0.565 | n.s. |
| DLSc | 0.005 | 2 | 171 | 0.012 | 0.194 | n.s. |
| DLSi | 0.021 | 1 | 86 | 0.012 | 1 | n.s. |

Supplementary Table 42: Statistics for Fig5Sup1d.

| Figure panel | Test | n (N mice) | Stat | p-value | Sig |
|---|---|---|---|---|---|
| a | Pearson's r | 22 (4) | -0.201 | 0.106 | n.s. |
| b | Pearson's r | 66 (4) | -0.312 | 0.011 | * |
| c | Wilcoxon (Low) | 36 (4) | 177 | 0.013 | # |
| | Wilcoxon (High) | 30 (4) | 168 | 0.191 | n.s. |
| d | Pearson's r | 66 (4) | 0.009 | 0.939 | n.s. |
| e | Pearson's r | 22 (4) | -0.216 | 0.345 | n.s. |
| f | Pearson's r | 23 (4) | 0.165 | 0.452 | n.s. |
| g | Pearson's r | 69 (4) | -0.143 | 0.241 | n.s. |

Supplementary Table 43: Statistics for Fig6Sup1.
